# Resource-access breadth marks protective strains against toxigenic *Clostridioides difficile*

**DOI:** 10.64898/2026.10.02.756234

**Authors:** Vishwas Mishra, Kevin Sia, Chloe Lindberg, Sneh Sharma, Michelle Wu, Ana Djukovic, Heesoo Jeong, Mini Kamboj, Qiwen Dong, Joao B. Xavier

**Author notes:** Centro de Investigación Príncipe Felipe (CIPF), Valencia, Spain.

## Abstract

Antibiotic disruption of gut microbiota creates opportunities for toxigenic *Clostridioides difficile*. One strategy is to occupy this niche with a non-toxigenic or weakly virulent strain, but even strains that cause little disease may fail to compete during pathogen challenge. We screened 21 sequence type 1 (ST1) isolates for their ability to protect mice co-colonized with toxigenic VPI10463, separating disease potential from protection. Some strains, such as ST1-75, caused little disease and protected, whereas others, such as ST1-68, caused little disease but failed to protect. Metabolomics showed that ST1-75 depleted more proline than ST1-68, whereas another protective strain, ST1-6, had a different metabolic profile; both protective strains nevertheless responded to many substrates. Substrate-utilization profiling revealed that protection across the 21 isolates correlated with amino-acid overlap with VPI10463, but not overall nutrient overlap or shared genome-encoded protein families. Relative amino-acid access explained variation beyond mono-colonization disease potential, and a consumer-resource model showed how nutrient overlap together with additional resource access can support persistence during competition. Additional experiments support that ST1-mediated protection occurs even in the absence of mature B and T cells. These findings distinguish low disease potential from protection, link protection to strain competition, and identify relative resource access as an informative phenotype for prioritizing strains for biotherapeutic testing.

**Importance:** *Clostridioides difficile* causes antibiotic-associated intestinal disease, but beneficial *C. difficile* strains could prevent this disease by competing with harmful strains. Selecting such strains requires more than identifying strains that cause little disease on their own: they must also compete effectively against a virulent one. We show that related *C. difficile* strains vary in their protective ability, and we linked this to their capacity to access amino acids and other resources used by the pathogen. Our findings highlight resource access as a key trait for selecting candidate strains and offer an ecological framework for developing live biotherapeutics against *C. difficile* disease.

## Introduction

Colonization resistance is a major defense against enteric infection. Resident microbes can restrict pathogens by consuming nutrients, producing inhibitory compounds, modifying bile-acid pools, occupying physical niches, and shaping host immunity (Buffie and Pamer 2013; Buffie et al. 2015; Round and Mazmanian 2009; Sorbara and Pamer 2019). Antibiotics weaken these barriers, allowing toxigenic *Clostridioides difficile* to expand, cause colitis, and recur after treatment (Lessa et al. 2015; McDonald et al. 2018; Theriot et al. 2014). Restoring colonization resistance is therefore an important therapeutic goal.

One strategy is to occupy the pathogen’s niche with a non-toxigenic or weakly virulent *C. difficile* strain. Asymptomatic colonization has been associated with a lower risk of subsequent *C. difficile*-associated diarrhea, and non-toxigenic strains can protect animals and patients from disease caused by toxigenic strains (Borriello and Barclay 1985; Gerding et al. 2015; Merrigan et al. 2003; Sambol et al. 2002; Shim et al. 1998). Weakly virulent toxigenic strains can also protect mice against more virulent strains (Etienne-Mesmin et al. 2018; Leslie et al. 2021). In patients with mixed-strain infections, the presence of non-toxigenic strains has been associated with less severe disease (Rodrigues-Jesus et al. 2026). This approach exploits the ecological similarity between the therapeutic strain and the pathogen. But it also creates a selection problem: a candidate must cause little disease on its own and remain competitively fit when a toxigenic strain enters the same disrupted ecosystem.

Work on gut communities provides an ecological framework for this selection problem. Nutrient overlap between bacteria in the resident gut microbiota and invading pathogens can promote colonization resistance through nutrient blocking (Spragge et al. 2023). Access to resources outside a resident’s niche, by contrast, can enable pathogen establishment, even when the resident and pathogen are closely related, such as when both are strains of *Escherichia coli* (Bakkeren et al. 2025). These findings suggest that selecting a protective strain requires considering both its overlap with the pathogen and the resource-access differences that support its own persistence.

Previous work identified ST1-75, a naturally low-disease clinical isolate belonging to *C. difficile* multilocus sequence type 1 (ST1), as a valuable starting point for this investigation. The ST1 lineage of *C. difficile* also includes epidemic toxigenic strains, and ST1-75 protected mice during simultaneous challenge with R20291, a toxigenic ST1 strain, or VPI10463. That study also showed that ST1-75 depletes amino acids faster than R20291, and amino-acid supplementation removes its competitive advantage (Dong et al. 2025). These findings established a mechanism for one successful protective strain but left open whether that mechanism could guide the selection of other protective strains or whether different protective strains use the same metabolic program.

Resource utilization varies among *C. difficile* strains, is an important determinant of their fitness, and could provide a basis for a general selection principle for avirulent strains to protect against a virulent challenger. Trehalose utilization can alter the behavior of epidemic *C. difficile* lineages (Collins et al. 2018), and amino-acid availability can promote pathogen expansion in dysbiotic communities (Batteaglioli et al. 2018). *C. difficile* nutrient use varies across gut environments (Jenior et al. 2017), while commensals can alter nutrient availability, metabolism, toxin production, and disease (Girinathan et al. 2021). Suppression can depend on Stickland-fermenting community members (Tian et al. 2025), although the importance of individual pathways varies across antibiotic-disrupted ecosystems (Huang et al. 2025). Here we asked which closely related *C. difficile* strains combine low disease potential with resource profiles that support competition against the pathogen, and which resource classes matter in this setting.

Closely related *C. difficile* isolates provide a system for answering this question. Isolates can differ in disease severity, colonization, substrate-supported activity, regulation, and accessory gene content even within the same lineage (Dong et al. 2023; Lewis et al. 2017; Nhu et al. 2025; Norsigian et al. 2022). Here, we used related clinical isolates from the ST1 lineage to ask why strains that cause little disease differ in protection (Miles-Jay et al. 2021). We reproduced ST1-75 protection in our laboratory and tracked its increase in relative frequency during direct competition with VPI10463, the toxigenic reference strain used for challenge in our mouse model. A strain screen identified ST1-68 as a closely related isolate that also caused little disease but provided weak protection. We then compared the strains metabolically and tested whether genomic and measured nutrient overlap, together with resource access relative to VPI10463, explained the variation in protection across the ST1 panel. A minimal consumer-resource model showed how the measured substrate-response patterns could support strain persistence, while immune and residual-microbiota experiments examined other biological differences. The resulting study separates disease potential from protective efficacy and proposes that relative resource access is a phenotype for prioritizing avirulent strains for ecological protection against *C. difficile* disease.

## Results

### ST1-75 protects and increases in relative frequency during VPI10463 challenge

Dong et al. showed that ST1-75 protects mice during simultaneous challenge with VPI10463 and outcompetes this toxigenic strain (Dong et al. 2025). We reproduced the protection phenotype in our laboratory at Memorial Sloan Kettering Cancer Center across seven experimental batches. Mice inoculated with a 1:1 mixture of ST1-75+VPI10463 maintained more weight than VPI10463-only controls in the same experiments (**Figure 1A**) and had markedly higher survival (**Figure 1B**; stratified log-rank p = 0.00435). To measure relative competitive fitness from a lower initial ST1-75 proportion, we used a different set of strain-specific quantitative PCR (qPCR) primers to track strain composition in mice co-inoculated with a 1:5 mixture of ST1-75:VPI10463. The mixed-infection mice lost weight after inoculation, but all remained above the VPI10463-alone trajectory at days 2 and 3, providing evidence of partial protection (**Figure 1C**). At 0.5 day after inoculation, ST1-75 was already at a mean fraction of 60%, rising to 84% on day 1, and each mouse exceeded 88% ST1-75 on day 3 (**Figure 1D**). This increase in relative frequency demonstrated a competitive advantage for ST1-75 against VPI10463. Protection and competitive enrichment were therefore reproduced in a second laboratory despite differences in inoculum ratio, sampling schedule, and qPCR assay.

**Figure 1.**
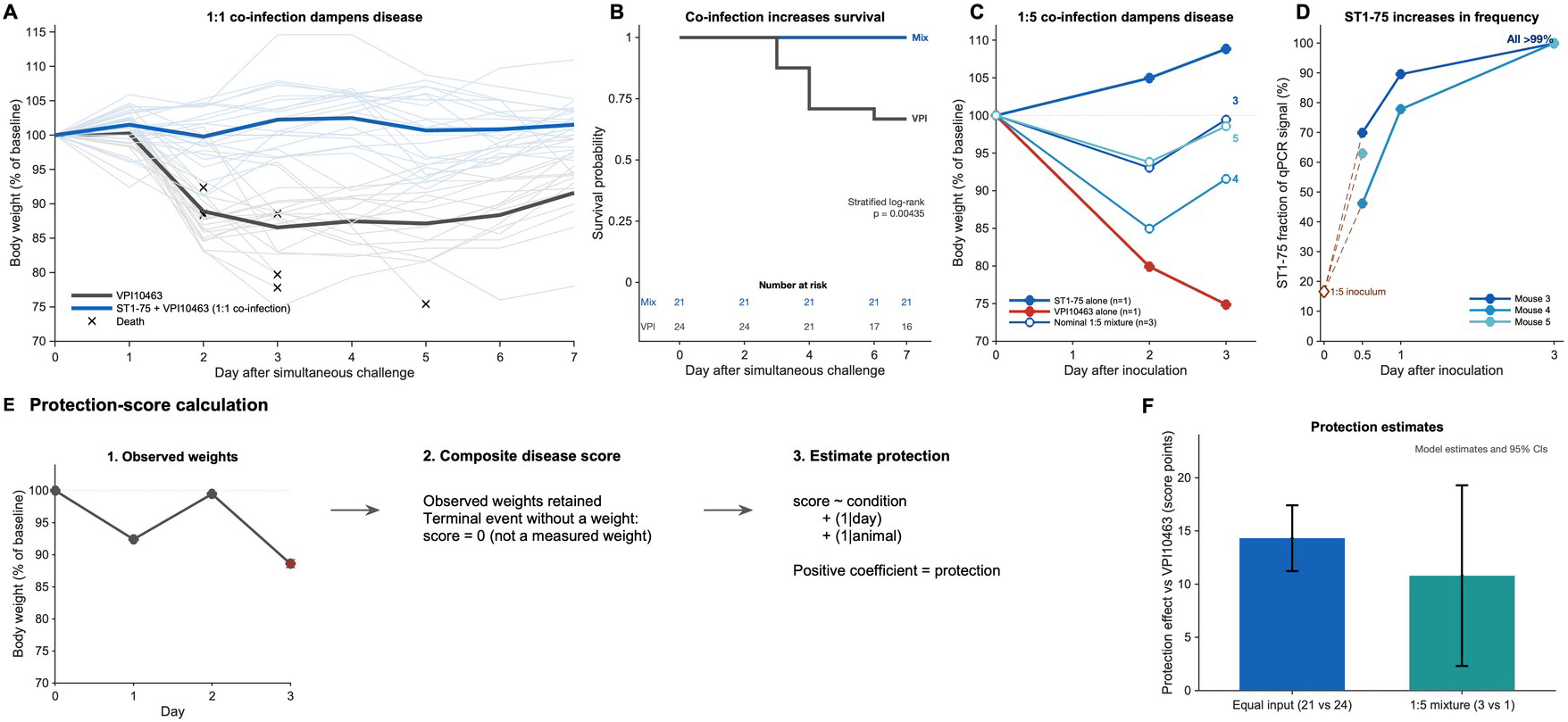
ST1-75 protects and increases in relative frequency during direct competition. (A) Relative body weight trajectories for mice co-infected with 1:1 ST1-75+VPI10463 and mono-infected VPI10463-only controls, compiled across 7 experimental batches. Thein lines show individual mice and bold lines show daily means calculated by giving each experiment equal weight. Crosses mark final weights before death. (B) Kaplan–Meier survival curves showing the numbers at risk (p = 0.00435). (C) Relative-weight trajectories in the 1:5 ST1-75+VPI10463 competition experiment shown for one ST1-75-alone mouse, one VPI10463-alone mouse, and three mixed-infection mice. Mixed-infection trajectories are labeled by mouse number and colored consistently with panel D. (D) ST1-75 relative qPCR fractions for the 3 mixed-infection mice (labeled mice 3, 4, and 5); the open diamond marks the 1:5 inoculum prepared, with dashed lines guiding the eye to the first observations, and mouse 5 lacks a day-1 value because no fecal pellet could be collected. (E) Protection-score calculation: Deaths contributed zero scores; a mixed-effects model fits condition as a fixed effect with random intercepts for day and animal identity, and its strain-level condition effect is the protection score; positive values indicate protection relative to VPI10463 alone. (F) Protection scores from the 1:1 and 1:5 experiments. Bars show model estimates, and whiskers show model-derived 85% confidence intervals.

Reproducing this phenotype in our laboratory provided the basis for developing a quantitative protection score from the longitudinal disease courses to compare candidate protective strains. Deaths contributed zero scores, allowing the protection score to incorporate terminal events (**Figure 1E**). Then, we used a mixed-effects model to estimate the condition effect relative to VPI10463 while accounting for repeated observations by day and animal; this strain-level effect is the protection score. Protection therefore denotes improved disease outcomes relative to VPI10463 alone, not necessarily absence of disease or dominance of the protective strain. ST1-75+VPI10463 produced a protection score of 14.31 points in the 1:1 inocula (95% CI, 11.23 to 17.40; 21 mixed-infection versus 24 VPI10463 mice) and 10.79 points in the 1:5 inocula (model-derived 95% CI, 2.29 to 19.28; three mixed-infection versus one VPI10463 mouse; **Figure 1F**). The wider confidence interval in the 1:5 experiment reflects its smaller sample size.

### Screen in mice separates disease potential from protection

We applied the protection score introduced with ST1-75 (**Figure 1E**) to a screen of 21 ST1 isolates. We used previously published data acquired from mono-colonization experiments (Dong et al. 2023) to measure disease potential by comparing each isolate with uninfected controls. We next conducted new 1:1 co-infection experiments with VPI10463 to measure protection by comparing each ST1+VPI10463 with VPI10463 alone. Higher disease scores indicate worse outcomes during mono-colonization; larger positive protection scores indicate better outcomes during toxigenic challenge, and the 95% confidence intervals quantified the uncertainty in those measurements. We found that across the 21 ST1 isolates, mono-colonization disease and co-infection protection were inversely associated (Spearman ρ = −0.721, p = 0.000337; **Figure 2C**). ST1-75 ranked second in protection and combined a nearnull disease effect (−0.55; 95% CI, −2.55 to 1.45) with strong protection (16.31; 95% CI, 10.23 to 17.40). ST1-6 had the largest protection estimate (16.31; 95% CI, 10.12 to 22.51) and a similarly near-null disease effect (−1.24; 95% CI, −4.55 to 2.07). By contrast, ST1-68 ranked last in protection despite a near-null disease effect (0.97; 95% CI, −2.34 to 4.28), with a protection estimate of −1.47 (95% CI, −8.05 to 5.12). ST1-49 showed the converse phenotype: it ranked sixth in protection (11.17; 95% CI, 4.97 to 17.37) despite a higher mono-colonization disease estimate (3.23; 95% CI, −0.54 to 7.00) (**Figure 2A-C**). Therefore, low disease potential did not ensure protection, and protection alone did not make a strain suitable as a low-disease therapeutic.

**Figure 2.**
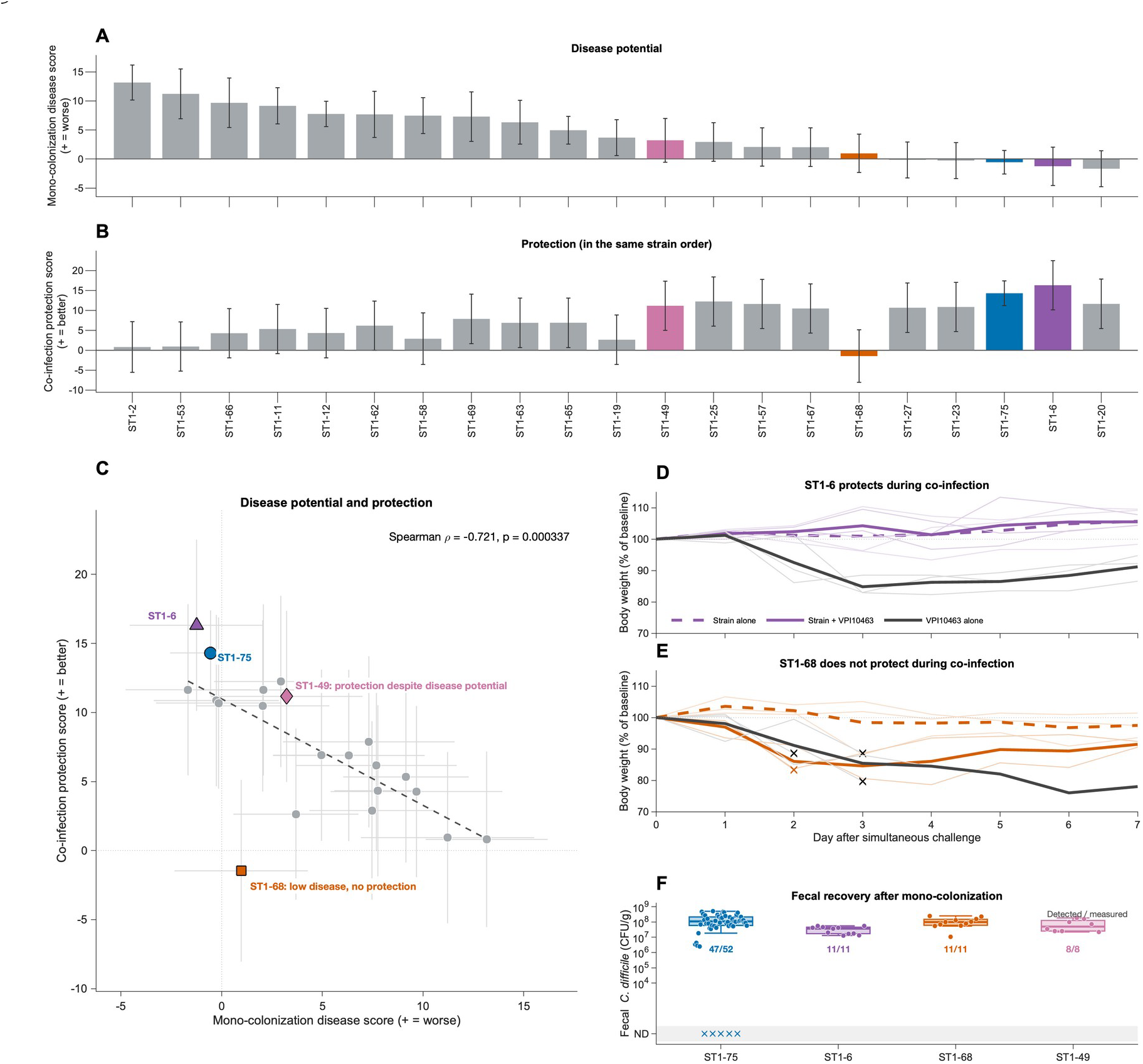
Low disease potential does not ensure protection during toxigenic challenge. (A) Mono-colonization disease scores for all 21 ST1 strains, ranked from greatest to least. Positive scores indicate worse outcomes than uninfected controls. (B) Co-infection protection scores shown in the identical strain order. Positive scores indicate better outcomes than VPI10463-only controls. Bars show model estimates and whiskers show model-derived 95% confidence intervals; horizontal lines mark the null reference. The strains discussed in the text, ST1-6, ST1-75, ST1-49, and ST1-68, are colored across panels. (C) Mono-colonization disease versus co-infection protection across the same 21 strains. Horizontal and vertical bars show 95% confidence intervals; the dashed line shows the ordinary least-squares fit, whereas the reported statistic uses Spearman correlation. ST1-68 caused little disease but provided weak protection, whereas ST1-49 reduced disease relative to VPI10463 alone despite a higher mono-colonization disease estimate. ST1-75 and ST1-6 combined low disease with strong protection. (D,E) Weight trajectories from the co-infection-screen experiments for ST1-6 (D) and ST1-68 (E), each shown with the VPI10463 controls run in the same experiment. Each experiment contained the ST1 strain alone, ST1+VPI10463, and VPI10463 alone. Thein lines show individual mice; dashed colored lines show strain-alone means, solid colored lines show ST1+VPI10463 means, and solid grey lines show VPI10463 means among observed animals. The ST1-6 experiment contained 3 mice per group and no terminal events. The ST1-68 experiment contained 3 strain-alone mice, 3 ST1-68+VPI10463 mice, and 4 VPI10463 mice; crosses mark the final observed weights before death in one ST1-68+VPI10463 mouse and three VPI10463-only mice. No terminal weight was imputed. (F) Fecal CFU/g measured on day 6 or 7 after mono-colonization with ST1-75, ST1-6, ST1-68, or ST1-49, according to the experiment-specific sampling schedule. Samples represent mice with an available late fecal measurement. Boxes span the interquartile range among positive log10 CFU/g measurements, center lines show medians, whiskers extend to the most extreme observations within 1.5 interquartile ranges, and circles show individual positive measurements. Crosses in the separate non-detect band preserve recorded zero counts without assigning a pseudocount. Labels give detected/measured samples: 47/52 for ST1-75, 11/11 for ST1-6, 11/11 for ST1-68, and 8/8 for ST1-49.

The time series from infected mice illustrate the differences in protective phenotype well (**Figure 2D,E**). Mice colonized with ST1-6+VPI10463 maintained higher weights than the VPI10463 controls, consistent with the positive protection score. ST1-68 alone also produced little disease, but ST1-68+VPI10463-colonized mice developed substantial weight loss, and one reached the composite endpoint of death or humane-endpoint removal (stratified log-rank p = 0.453; Benjamini-Hochberg-adjusted q = 0.480 across the strain-panel comparisons). The screen therefore identified ST1-6 and ST1-68 as low-disease strains with very different protection phenotypes.

To determine whether ST1-68 failed to protect because it could not colonize the antibiotictreated mice, we compared late fecal loads collected on days 6–7 after mono-colonization (**Figure 2F**). ST1-75 was detected in 47 of 52 measured samples, ST1-6 in 11 of 11, ST1-68 in 11 of 11, and ST1-49 in 8 of 8. Positive ST1-68 burdens ranged from 1.04×10^7^ to 2.50×10^8^ colony-forming units per gram of feces (CFU/g), within the range observed for the other highlighted strains. Each strain could therefore establish during mono-colonization despite its different protection phenotype. Across all 21 isolates, we recovered every ST1 strain from at least one mouse (**Supplementary Figure S1A,B**), consistent with previous findings that these ST1 strains colonize antibiotic-treated mice (Dong et al. 2023).

In co-colonized mice, plating measures total *C. difficile* because colonies cannot be assigned to the ST1 strain or VPI10463. Nonetheless, this combined burden did not correlate with protection across the strain panel (**Supplementary Figure S1C,D**). ST1-68 established during mono-colonization, but these total co-colonization counts do not establish its persistence alongside VPI10463. Differences in total fecal burden did not explain differences in protection against disease.

### Protective strains have distinct metabolic phenotypes

ST1-75 and ST1-68 provided a direct comparison: both produced little disease on their own, yet only ST1-75 protected during toxigenic challenge. We used metabolic profiling to ask whether this divergence was accompanied by differences in intracellular bacterial metabolites and in the compounds the strains consumed from or released into the medium.

We used gas chromatography–mass spectrometry (GC-MS) and an established derivatization protocol that detects several amino acids and tricarboxylic acid (TCA)-cycle intermediates (e.g., (Xavier 2024)) to examine intracellular metabolites across three matched experimental batches after six hours of growth in chemically defined *C. difficile* minimal medium (CDMM). The five strains included ST1-75 and ST1-68, ST1-6 as another strongly protective isolate, VPI10463 as the present challenge strain, and R20291 as the toxigenic model strain used in the previous ST1-75 study (Dong et al. 2025). Partial least-squares discriminant analysis (PLS-DA) placed ST1-75 and ST1-68 in different regions, while ST1-6 overlapped substantially with ST1-68 (**Figure 3A**). No individual intracellular metabolite distinguished ST1-75 from ST1-68 after false-discovery correction.

**Figure 3.**
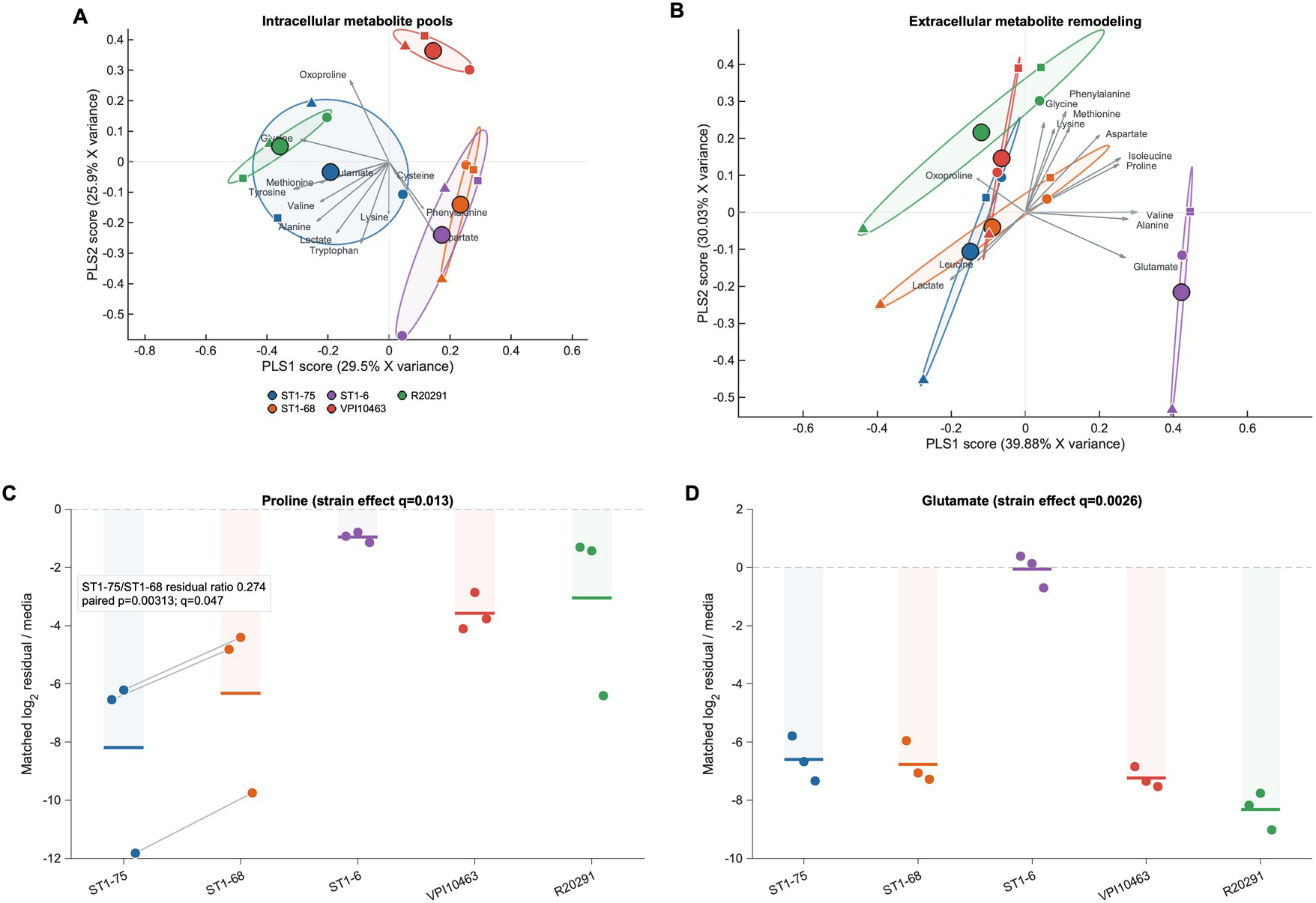
Intracellular and extracellular metabolite profiles across strains. (A) Partial least-squares discriminant analysis (PLS-DA) of intracellular metabolite pools after subtraction of the batch-matched blank medium. The analysis includes 13 of 18 measured metabolites whose blank-corrected grand mean was above background at Benjamini-Hochberg q < 0.05. Small symbols show individual samples, symbol shape denotes matched batch, large circles show strain centroids, arrows show scaled metabolite loadings, and colored ellipses outline the spread of the three observed samples within each strain. (B) PLS-DA of the 13 continuously quantified extracellular batch-matched log2 signals relative to the corresponding uninoculated medium. Symbols, centroids, loading arrows, and ellipses use the same conventions as in A. The first two components explain 39.88% and 30.03% of extracellular X variance, respectively. The two compartment-specific projections were fitted independently. (C) Extracellular proline signals for the 5 strains, expressed as batch-matched log_2_ ratios relative to uninoculated medium; zero indicates the medium level. Lines connect matched ST1-75/ST1-68 batches. The ST1-75/ST1-68 geometric residual-signal ratio was 0.274 (paired p = 0.00313, Benjamini-Hochberg q = 0.04698 across 15 nonconstant metabolites; exact two-sided sign-flip p = 0.25). The 5-strain global test gave q = 0.0130. (D) Extracellular glutamate signals on the same scale; the five-strain global test gave q = 0.0026. Bars in C and D show group means; pale fills extend from zero to each mean. Each group contains 3 matched experimental batches.

PLS-DA of extracellular signals relative to uninoculated medium processed in the same batch organized the strains differently (**Figure 3B**). ST1-6 separated strongly along the first component, whereas ST1-75, ST1-68, VPI10463, and R20291 overlapped substantially. Intracellular pools and extracellular remodeling therefore distinguished different relationships among the strains.

At the feature level, proline and glutamate were the only continuously quantified metabolites with global strain effects after false-discovery correction (**Figure 3C,D**). Decreases relative to blank CDMM medium indicate net depletion from the extracellular pool (metabolite consumption by the strain), whereas increases indicate net accumulation (metabolite release by the strain). ST1-75 left less residual proline than ST1-68 in each matched batch (**Figure 3C**), identifying a faster consumption rate.

ST1-6 consumed less proline and glutamate than ST1-75 (**Figure 3C,D**). The two protective strains therefore did not share a pattern of extracellular metabolite consumption/release, prompting us to ask whether protection was associated with access to a wider range of resources.

### Nutrient overlap and relative amino-acid access mark protection

The protection differences between ST1-75 and ST1-68 should ultimately be driven by genetic differences. Genome-wide comparison distinguished these closely related strains at 45 nucleotide positions across 3.50 million jointly readable bases outside inferred recombination regions (**Figure 4A**). Across all 21 strains, phylogenetic distance was modestly associated with pairwise differences in mono-colonization disease (ρ = 0.183, p = 0.025) and co-infection protection (ρ = 0.238, p = 0.008).

**Figure 4.**
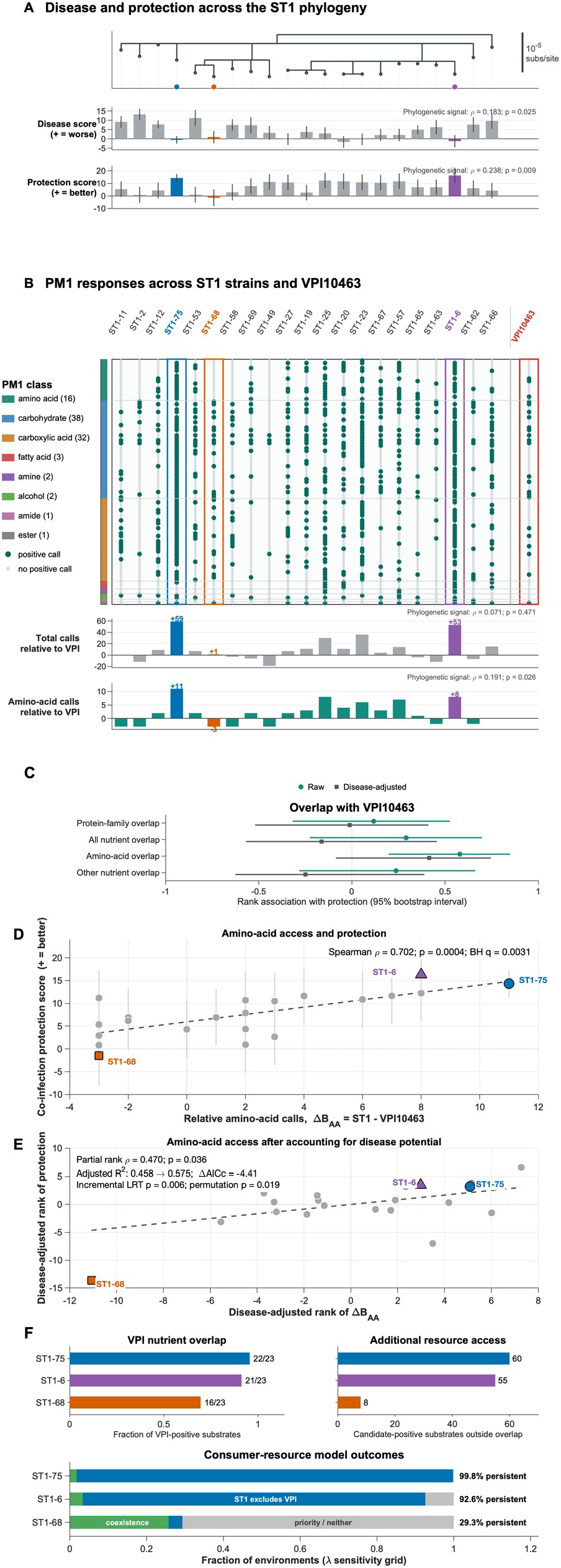
Nutrient overlap and relative amino-acid access in protective-strain selection. (A) VPI10463-rooted, recombination-filtered whole-genome phylogeny of 21 ST1 strains, with VPI10463 removed from the displayed tree. Internal branches with ultrafast bootstrap support below 95% or SH-aLRT support below 80% are collapsed for display; phylogenetic tests use the full fitted tree. Aligned bar tracks show mono-colonization disease and co-infection protection effects. Whiskers show model-derived 95% confidence intervals. Annotations report two-sided Mantel-style Spearman label-permutation tests between patristic distance and pairwise trait divergence. (B) Complete binary PM1 substrate growth calls in the identical strain order. Rows are grouped by chemical class. Large teal circles indicate positive growth calls. The aligned tracks show total relative access, ΔB = B_ST1_ - 23, and amino-acid relative access, ΔB_ST1_ = B_ST1,AA_ - 3. Track annotations report the corresponding phylogenetic tests. (C) Raw and disease-adjusted rank associations between protection and VPI10463-centered protein-family or nutrient overlap across 21 strains. Circles and squares show raw and adjusted coefficients, respectively; horizontal lines show percentile 95% intervals from 5,000 strain bootstraps. The vertical dotted line marks zero. Nutrient overlap is the fraction of VPI10463-positive substrates also positive for the candidate, with denominators of 23 overall, 3 amino-acid-class substrates, and 20 remaining substrates. Protein-family overlap uses the 2,983 distinct VPI10463 global protein families (PGFams). (D) ΔB_AA_ versus the protection score across the 21 ST1 strains. Vertical intervals are 95% score confidence intervals; the dashed line shows the ordinary least-squares fit, whereas the reported statistic uses Spearman correlation. The displayed q-value is from the eight-class comparison in **Supplementary Figure S2D**. (E) Rank association between ΔB_AA_ and protection after both quantities were residualized for mono-colonization disease rank. The annotation reports the partial-rank association and the improvement obtained by adding ΔB_AA_ to the disease-only model; adjusted R^2^ is from ordinary least squares, while ΔAICc and the likelihood-ratio test are from generalized least squares using the covariance among protection-score estimates. (F) Upper: VPI10463 nutrient overlap and additional candidate resource access outside that overlap for ST1-75, ST1-6, and ST1-68. Lower: consumer-resource outcomes for the same complete all-resource call patterns across 501 values of the aggregate resource-limitation parameter λ. ST1 persistence combines coexistence and ST1 exclusion of VPI10463. Grey combines priority effects, for which the outcome depends on which strain establishes first, and parameter values for which neither strain is feasible. ST1-75, ST1-6, and ST1-68 are highlighted across panels.

We first asked whether genome-encoded protein-family overlap provided a correlate of protection. Following the pathogen-centered measure developed previously (Spragge et al. 2023), we calculated the fraction of VPI10463-encoded protein families also represented in each candidate’s genome. The 21 ST1 strains covered 87.2–88.9% of the 2,983 assigned VPI10463 families, but overlap was not detectably associated with protection, either before or after accounting for disease potential (ρ = 0.118 and partial-rank ρ = −0.013, respectively; **Figure 4C**).

Because ST1-75 and ST1-6 did not share one extracellular consumption profile, we asked whether protection instead correlated with access to a wider range of resources. We used BIOLOG PM1 plates to expose each strain separately to 95 defined carbon-source environments and converted the recorded optical-density-at-600-nm (OD600) growth curves into binary positive or negative substrate-response calls. We refer to the number of positive calls as response breadth. Closely related strains nevertheless responded to different sets of substrates (**Figure 4B**; **Supplementary Figure S2A**). ST1-75 and ST1-6 showed detectable growth on 82 and 76 substrates, compared with 24 for ST1-68 and 23 for VPI10463. Within the amino-acid class, ST1-75 and ST1-6 responded to 14 and 11 of 16 substrates, whereas ST1-68 and VPI10463 responded to 0 and 3.

We defined nutrient overlap as the fraction of VPI10463-positive substrates also positive for the candidate. Overall overlap was not detectably associated with protection (ρ = 0.291). Amino-acid overlap showed a positive association (ρ = 0.579, Benjamini–Hochberg q = 0.0355 across six overlap tests), whereas overlap across the remaining substrates did not (**Figure 4C**). The disease-adjusted amino-acid estimate was less certain (partial-rank ρ = 0.415, p = 0.069, q = 0.208; 95% bootstrap interval, −0.086 to 0.746).

Overlap describes the resources on which strains can compete, but does not capture access beyond those resources. We therefore also calculated relative access, ΔB, as candidate total breadth minus VPI10463 breadth, and relative amino-acid access, ΔB_AA_, as candidate amino-acid breadth minus VPI10463 amino-acid breadth. ST1-75 had ΔB = +58 and ΔB_AA_ = +11; ST1-6 had +53 and +8; and ST1-68 had +1 and −3, respectively (**Figure 4B**). Pairwise differences in total relative access were not associated with phylogenetic distance (ρ = 0.071, p = 0.471), whereas amino-acid access showed a modest association (ρ = 0.191, p = 0.026). The phylogenetic associations with disease, protection, and amino-acid access were stable to outgroup choice but were not detected before recombination filtering.

Previous work on ST1-75 and its faster proline consumption led us to examine amino-acid access specifically. We also asked whether this phenotype explained protection beyond the differences associated with mono-colonization disease potential. Among eight chemical classes,amino-acid access showed the strongest association with protection; fatty-acid and amine breadth also passed false-discovery correction but were based on only 3 and 2 PM1 substrates, respectively (**Supplementary Figure S2D**). Across the 21 ST1 strains, ΔB_AA_ correlated with protection (Spearman ρ = 0.702, p = 0.00039, Benjamini–Hochberg q = 0.0031; 5,000-resample 95% interval, 0.292 to 0.926; **Figure 4D**; **Supplementary Figure S2B**). Across the remaining 79 non-amino-acid environments, the raw association was weaker (ρ = 0.386, p = 0.0841) and little association remained after disease adjustment (partial-rank ρ = 0.082, p = 0.730; **Supplementary Figure S2B**).

To make this comparison, we removed the rank relationship with mono-colonization disease from both amino-acid access and protection, then tested the association between the remaining variation. The amino-acid relationship remained detectable (partial-rank ρ = 0.470, p = 0.0365; **Figure 4E**). Adding ΔB_AA_ to the disease-only model increased adjusted R^2^ from 0.458 to 0.575 and improved covariance-aware generalized least-squares fit (difference in small-sample corrected Akaike information criterion, ΔAICc = −4.41; likelihood-ratio p = 0.0062; residual-permutation p = 0.0188). Relative amino-acid access remained associated with protection in a model accounting for disease potential, phylogenetic relatedness, and uncertainty in protection scores (parametric-bootstrap p = 0.013). In a joint model containing disease, amino-acid breadth, and non-amino-acid breadth, the amino-acid term remained supported (residual-permutation p = 0.0260), whereas the non-amino-acid term did not (p = 0.412). Removing any one of the 16 amino-acid-class substrates left the raw correlation between 0.676 and 0.719 and the disease-adjusted correlation between 0.442 and 0.510, showing that no single substrate carried the class-level association (**Supplementary Figure S2C**). Selecting one PM1 plate per strain preserved the association across all available plate combinations (raw ρ = 0.675–0.718; disease-adjusted ρ = 0.437–0.480). The disease-adjusted correlation strengthened without ST1-49 (ρ = 0.621) but weakened without ST1-68 (ρ = 0.131). Adding amino-acid access improved explanatory fit but not held-out prediction: leave-one-strain-out root-mean-square error (RMSE) was 3.679 for disease alone and 3.680 after adding amino-acid access.

ST1-75 and ST1-6 combined high nutrient overlap with substantial additional resource access, defined as candidate-positive substrates outside the overlap (**Figure 4F**). ST1-75 overlapped with 22 of the 23 VPI10463-positive substrates and accessed 60 additional substrates; ST1-6 overlapped with 21 and accessed 55 additional substrates. ST1-68 overlapped with 16 and accessed only eight additional substrates, while seven VPI10463-positive substrates lay outside its resource profile. The two assays agreed locally for proline: ST1-75 grew on proline and left less extracellular proline than ST1-68, which failed to grow on proline. This agreement did not extend to the substrate-level or whole-profile comparisons, indicating that the two assays captured different aspects of strain metabolism.

### Consumer-resource model links resource access to persistence during competition

We used a minimal consumer-resource model to ask how nutrient overlap and additional resource access across all 95 PM1 environments could affect persistence during toxigenic challenge. The model used the number of overlapping resources and each strain’s total access, with access outside the overlap derived from these counts. Across 501 values of an aggregate resource-limitation parameter, the ST1-75 and ST1-6 profiles were compatible with ST1 persistence at 99.8% and 92.6% of values, respectively, compared with 29.3% for ST1-68 (**Figure 4F**).

### ST1-75-colonized RAG1-deficcient mice maintain weight after challenge

To explore additional mechanisms that could be involved in the protection phenotype, we examined the response of ST1-75-colonized mice lacking mature adaptive immunity to VPI10463 challenge. ST1-75-colonized wild-type mice and mice deficient in recombination-activating gene 1 (RAG1), which lack mature B and T lymphocytes (Mombaerts et al. 1882), maintained their weight after VPI10463 challenge, whereas previously uncolonized wild-type controls developed pronounced weight loss (**Figure 5A**). Using the same mixed-effects model structure as in the mouse screen, fitted to observed weights, each ST1-75-colonized arm showed a positive protection effect relative to the previously uncolonized wild-type arm (**Figure 5B**). ST1-75-colonized mice therefore maintained their weight after challenge even in the absence of mature B and T cells.

**Figure 5.**
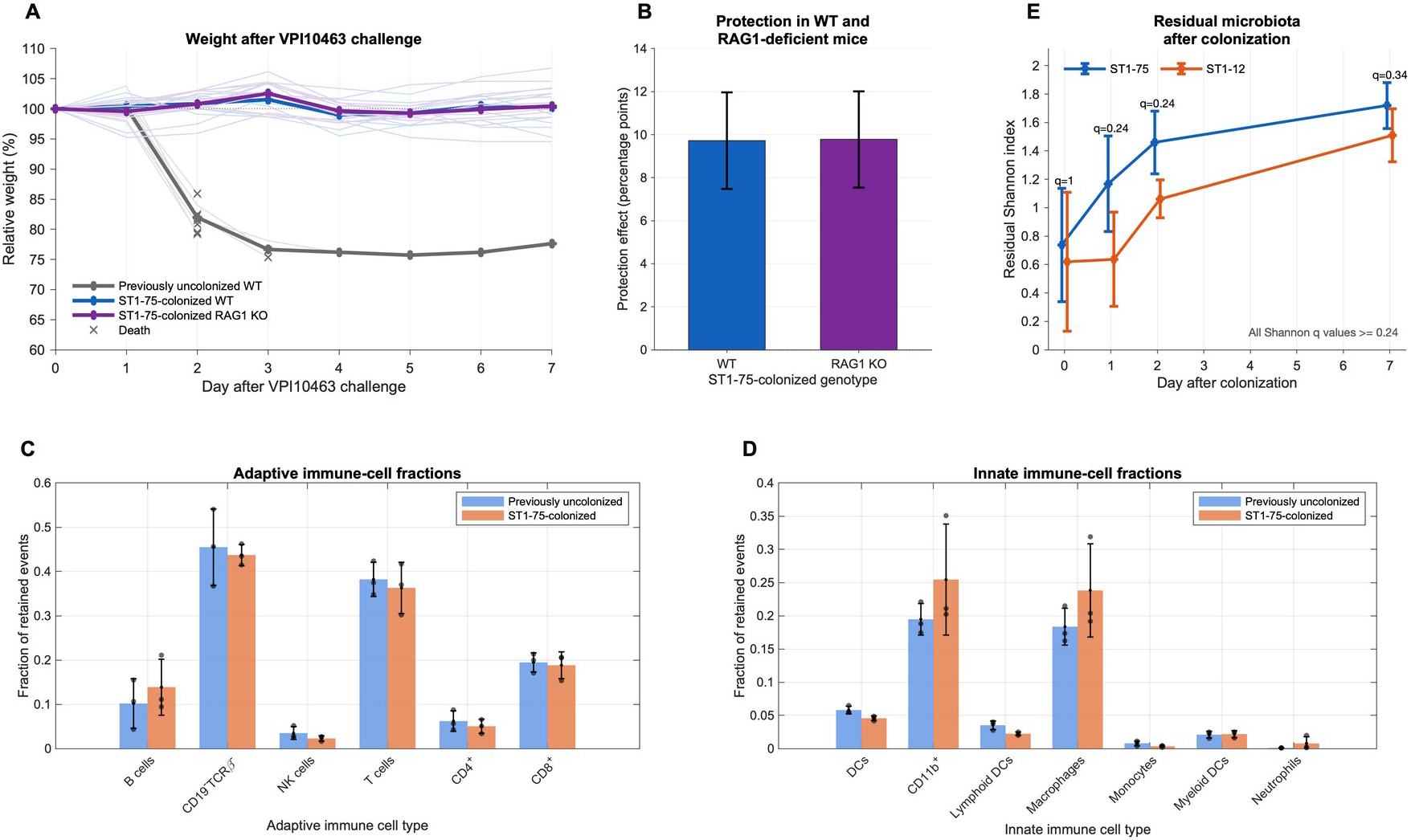
Adaptive immunity and residual microbiota. (A) Weight trajectories after VPI10463 challenge in previously uncolonized wild-type mice and in ST1-75-colonized wild-type or RAG1-deficient mice (n=10 mice per group). Thein lines show individual mice and bold lines show group means among observed animals. Crosses mark the death of the 9 previously uncolonized wild-type mice who succumbed to the infection. (B) Protection effects for the ST1-75-colonized wild-type and RAG1-deficient arms, each estimated relative to the previously uncolonized wild-type arm using the mixed-effects disease-course model: relative weight as outcome, condition as a fixed effect, and random intercepts for day and animal. Both bars represent ST1-75-colonized mice and use the same previously uncolonized wild-type reference. The x-axis labels identify the genotype of the colonized arm. Bars are model estimates and whiskers are 95% confidence intervals. (C,D) Adaptive- and innate-immune-cell fractions measured in a separate flow-cytometry experiment comparing previously uncolonized and ST1-75-colonized mice (3 mice per group). Bars show group means, points show individual mice, and error bars show standard deviations. Cell fractions were calculated relative to the total retained events for each mouse within each staining panel (see Methods). (E) Shannon diversity of the residual genus-level microbiota after removal of *Clostridioides* and renormalization of the remaining community in mice colonized with ST1-75 or the higher-disease comparator ST1-12. Points show group means and error bars show standard deviations; each strain and day contains 3–5 mice. Labels give Benjamini-Hochberg-adjusted q values. **Supplementary Figure S3A,B** shows the community-composition and genus-level comparisons.

In a separate flow-cytometry experiment, three previously uncolonized and three ST1-75-colonized mice showed only modest differences, without a dominant redistribution of the measured adaptive or innate immune-cell fractions (**Figure 5C,D**). We also compared residual microbiota between mice colonized with ST1-75 and mice colonized with ST1-12, a higher-disease isolate with a smaller protection estimate in the mouse screen. After removing *Clostridioides* and renormalizing the remaining genus-level community, none of the measured diversity, community-composition, or genus-abundance comparisons survived false-discovery correction (**Figure 5E; Supplementary Figure S3A,B**).

## Discussion

Protective-strain selection requires low disease potential together with competitive fitness, and relative resource access offers a phenotype for prioritizing candidates. ST1-75 reduced disease during simultaneous challenge with VPI10463, increased event-free survival, and produced a positive protection score. It also increased in relative frequency during competition. The enrichment experiment directly links protection, a host-level outcome, to relative competitive fitness between the strains. ST1-75-colonized mice also maintained their weight after challenge in the absence of mature B and T cells. Reproducing the previously reported protection phenotype (Dong et al. 2025) in a different laboratory, extending competition to a 1:5 inoculum, and developing a quantitative protection score established a basis for comparing candidate strains.

The screen of 21 ST1 isolates showed why this distinction matters for therapeutic selection. Disease potential and protection were strongly related, but low disease potential was insufficient to identify protective strains. ST1-68 had similarly low mono-colonization disease as ST1-75 but, unlike ST1-75, failed to protect against VPI10463-caused disease. Conversely, ST1-48 reduced disease relative to VPI10463 alone but had a higher disease score on its own; its positive protection score does not imply avirulence. Disease during co-infection can persist whether a virulent ST1 strain displaces VPI10463, coexists with it, or is displaced, so disease outcomes alone do not identify the competitive outcome. A candidate for therapeutic use must therefore satisfy two different requirements: low disease potential defines safety, while competitive fitness supports efficacy during toxigenic challenge.

Fecal burden measurements established that ST1-68 could colonize on its own. Total *C. difficile* burden during co-colonization did not follow the protection ranking, but did not distinguish the competing strains. Strain-specific measurements during ST1-68/VPI10463 co-infection would help determine whether weak protection accompanies poor persistence or persists despite successful colonization.

The comparison with ST1-68 supported an ecological explanation for ST1-75 efficacy. ST1-75 left less extracellular proline than this closely related strain, extending its previously described amino-acid phenotype (Dong et al. 2025). ST1-6, another strongly protective strain, did not reproduce the same extracellular profile. Protective strains can therefore occupy different realized metabolic states. Across the strain panel, amino-acid access relative to VPI10463 showed the strongest class-level relationship with protection and added explanatory information beyond mono-colonization disease. Breadth across the other 79 PM1 substrates did not show the same disease-adjusted relationship. Relative resource access is therefore an associated phenotype, not a universal explanation for protection; strains such as ST1-49 may reduce disease through other interactions. Genomic relatedness also showed modest associations with disease, protection, and amino-acid access. Small regulatory differences can nevertheless have substantial phenotypic effects: variation in flagellar-switch inverted repeats alters gene-expression heterogeneity and virulence within ST1 (Nhu et al. 2025). The consumer-resource model showed how overlap together with additional resource access, not amino-acid access alone, could expand the range of conditions compatible with ST1 persistence. Our consumer–resource model shares an invasion logic proposed previously (Bakkeren et al. 2025), but focuses on resource competition as the driver of candidate persistence. It does not address the potential contribution of antimicrobial interference to competition between strains.

These results connect nutrient blocking in communities (Spragge et al. 2023) to the selection of closely related strains for disease protection. In this ST1 panel, protein-family overlap varied narrowly and did not track protection. Its coarse representation of function may leave differences in regulation and effective nutrient use unresolved. Previous work showed that protein-family differences are informative for competition among *E. coli* strains (Bakkeren et al. 2025), so our result does not imply that genomic comparisons generally fail between close relatives. The substrate utilization profiling measured with BIOLOG plates provided greater phenotypic resolution here. The amino-acid overlap association and the complementary relative-access result also point to the importance of resource identity: competition depends on the pathogen, competitors, and environment, not simply the number of overlapping resources. These findings agree with the notion that amino-acid access is key in *C. difficile* ecology, but amino acids may be less central to protection against other pathogen species.

We should note that the panel-wide association is not a prediction rule. Its disease-adjusted estimate weakened when we removed ST1-68, and adding amino-acid access did not improve leave-one-strain-out error beyond disease alone. At this stage, relative amino-acid access can prioritize candidates for direct competition experiments but cannot substitute for measuring competitive fitness *in vivo*. It is also strain-specific: recalculate the same pairwise quantity for another toxigenic strain after measuring its resource-response profile. Future work should also investigate relative amino-acid access in newly selected strain–challenge pairs and combine absolute population measurements with time-resolved metabolite profiles. Nonetheless, our work highlights that protective-strain selection must consider both safety and the ability to compete with toxigenic *C. difficile*. By linking mouse phenotypes to competitive enrichment and pairwise resource access, this study expands a mechanism discovered in ST1-75 into a testable strategy for prioritizing live-biotherapeutic candidates. Extending this approach to other disrupted microbial ecosystems will require resolving both nutrient overlap and the access differences that support candidate persistence, and identifying which resources matter in each setting.

## Methods

### Bacterial strains and spore preparation

Clinical isolates belonging to *C. difficile* multilocus sequence type 1 (ST1; ribotype 027) were obtained through prospective molecular surveillance at Memorial Sloan Kettering Cancer Center between 2013 and 2017 (Miles-Jay et al. 2021). Twenty-one isolates were selected by multilocus sequence type for the strain panel. VPI10463 (ATCC 43255) was used as the toxigenic challenge strain in the mouse experiments; R20291 was included as an additional virulent strain in the GC-MS experiment.

To prepare spores, strains were streaked on brain heart infusion-supplemented (BHIS) agar containing taurocholic acid after overnight pre-reduction in an anaerobic chamber. After two days, a single colony was inoculated into 5 mL pre-reduced BHIS broth and cultured for 24 hours. This culture was transferred into 40 mL pre-reduced BHIS broth and incubated for 5–7 days. Cultures were centrifuged at 4,000 × g for 30 minutes at 4 °C. Pellets were resuspended in 1 mL cold water, stored at −20 °C for two days, thawed at room temperature, and washed five times with cold double-distilled water by centrifugation at 13,000 rpm for two minutes. The washed pellet was resuspended in 1 mL phosphate-buffered saline and heated at 70 °C for 20 minutes. Spore preparations were quantified from three serial dilutions plated anaerobically on pre-reduced BHIS agar containing taurocholic acid.

### Mouse infection models

Six- to eight-week-old conventionally housed female C57BL/6 mice were obtained from Jackson Laboratories, maintained under specific-pathogen-free conditions, and acclimated for one week. Mice were individually housed in autoclaved cages during infection experiments. Experiments were conducted under protocols approved by the Memorial Sloan Kettering Cancer Center Institutional Animal Care and Use Committee.

To disrupt colonization resistance, mice received metronidazole, neomycin, and vancomycin in drinking water, each at 0.25 g/L, for three days, followed by regular water for two days. Clindamycin (200 micrograms in 100 microliters) was administered intraperitoneally 24 hours before oral spore challenge. Mono-colonization experiments used approximately 200 spores of one ST1 strain, with uninfected mice processed as controls. Panel co-infection experiments used approximately 200 ST1 spores and approximately 200 VPI10463 spores; VPI10463-only mice served as controls for protection. Animals were monitored for body weight, stool consistency, clinical signs, death, and humane-endpoint removal. Death was defined as observed death or removal at a humane endpoint. Fecal samples were stored at −80 °C until analysis.

For the adaptive-immunity experiment, ST1-75 colonization used the same antibiotic pretreatment and approximately 200 ST1-75 spores. Before VPI10463 challenge, the antibiotic regimen was repeated and approximately 200 VPI10463 spores were administered. ST1-75-colonized wild-type and RAG1-deficient mice, which lack mature B and T lymphocytes (Mombaerts et al. 1992), were compared with previously uncolonized wild-type challenge controls; each group contained ten mice.

### ST1-75/VPI10463 mixed infection

Mixed infection experiments compared ST1-75 alone, VPI10463 alone, and a mixed infection designed with an approximately 1:5 ST1-75:VPI10463 inoculum target. Relative weights were recorded at days 0, 2, and 3 for one ST1-75-alone mouse, one VPI10463-alone mouse, and three mixed-infection mice (**Figure 1C**).

Fecal samples were collected 0.5, 1, and 3 days after infection. DNA was extracted using the QIAamp Fast DNA Stool Mini Kit (Qiagen, 51 604), with glass-bead disruption in InhibitEX buffer, heating at 95 °C for 7 minutes, proteinase K treatment, and elution in 50 microliters of nuclease-free water. Day-1 measurements shown in **Figure 1D** used fecal samples collected for CFU enumeration. No day-1 pellet could be collected from mouse 5 because of diarrhea.

Strain-specific qPCR targeted a 147-bp ST1-75 product with forward primer 5′-TGTTCCCATCCAGGCAAAAATC-3′ and reverse primer 5′-CTTCGTATATGGTGCGAGCG-3′, and a 168-bp VPI10463 product with forward primer 5′-GCGTTTCATGACCGTTCCAC-3′ and reverse primer 5′-TCCAGCGGCGACAAAGTTAT-3′. Reactions were run in technical triplicate on a QuantStudio 6 Flex in 10-microliter volumes containing 5 microliters of PowerUp SYBR Green Master Mix, 200 nM of each primer, and 2 microliters of template DNA. Cycling comprised 50 °C for 2 minutes, 95 °C for 2 minutes, and 40 cycles of 95 °C for 1 second and 60 °C for 30 seconds, followed by melting-curve analysis. Separate tenfold dilution series calibrated the two targets; amplification efficiencies were 93.46% for ST1-75 and 96.56% for VPI10463, with R^2^ = 0.9991 for both curves. Relative ST1-75 abundance was calculated as its mean copy estimate across technical replicates divided by the sum of the ST1-75 and VPI10463 mean copy estimates from the same sample, multiplied by 100.

Compared with our experiments, Dong et al.’s VPI10463 study was conducted at the University of Chicago, used 100 spores per strain at a 1:1 ratio with four mice per group, and reported day-1 qPCR (Dong et al. 2025). Their spore preparation included density-gradient purification. Their qPCR protocol used PowerSoil Pro or PowerFecal Pro DNA extraction, wild-type/common *)cdtR* primers, PowerTrack SYBR Green chemistry, a QuantStudio 6 Pro, and efficiency-based relative quantification. Our assay instead used separate copy calibration for the two strain-specific targets to calculate strain fractions.

### Longitudinal weight and survival analyses

Mono-colonization disease effects were estimated by comparing each ST1-alone condition with uninfected controls; positive effects indicate worse disease. Co-infection protection effects were estimated by comparing each ST1+VPI10463 condition with VPI10463-only controls; positive effects indicate improved disease course. The fitted strain-level effects define the disease and protection scores.

When relative weight was missing on a recorded day of death or humane-endpoint removal, the analysis assigned a score of zero; measured weights were retained. The model included condition as a fixed effect and random intercepts for day and individual mouse, with each mouse identified uniquely across experiments and conditions.

For the mono-colonization model, the relative-weight score was multiplied by −1 before fitting so positive coefficients represented worse disease than uninfected controls. Positive co-infection coefficients represented better disease-course scores than VPI10463-alone controls. Models included all experimental conditions represented in each comparison, and the reported correlation was restricted to the 21 ST1 strains measured in both screens.

Zero scores represent terminal events, not measured body weights. Raw weight trajectories show observed measurements only. For **Figure 1A**, faint lines show each animal’s observed weights. Bold lines are observed daily means computed first within each experiment and then averaged across experiments so that every experiment contributes equally. For animals with a terminal event, a cross marks the final measured weight on the trajectory.

Survival analyses treated observed deaths and humane-endpoint removals as events at the recorded day; animals without events were censored at their final recorded day. **Figure 1B** shows pooled Kaplan–Meier curves and numbers at risk for ST1-75+VPI10463 and contemporaneous VPI10463 controls. Inference used experiment-stratified log-rank tests comparing each ST1 condition with contemporaneous VPI10463 controls, summing observed-minus-expected totals and variances across strata. The ST1-75 comparison in **Figure 1** was reported as a single comparison without multiple-testing correction. Benjamini-Hochberg correction was applied across the panel-wide survival comparisons.

### Fecal burden

Fecal *C. difficile* burden was quantified as colony-forming units per gram of feces (CFU/g) by plating on selective agar (Dong et al. 2023). For each experiment, the early sampling window was day 1 or 2 and the late sampling window was day 6 or 7, following the experiment-specific collection schedule. Strain-alone measurements included both the dedicated mono-colonization experiments and the strain-alone control arms run alongside co-infection experiments. Measurements from candidate+VPI10463 groups were reported as total *C. difficile* CFU.

Statistical analyses used positive log10-transformed CFU/g measurements; recorded zeros were reported separately as non-detects without assigning a pseudocount, and missing measurements were excluded. For each sampling window, differences among strain conditions were tested by likelihood-ratio comparison of maximum-likelihood linear mixed-effects models with and without strain as a fixed effect. Both models included a random intercept for experiment; strain-alone models also included a fixed effect distinguishing measurements from the mono-colonization screen and strain-alone controls in the co-infection screen. Benjamini– Hochberg correction was applied across the four global comparisons (two sampling windows in each colonization context). Associations with disease or protection scores were assessed across the 21 ST1 strains using two-sided Spearman correlations. For each strain and sampling window, burden was summarized as the median of the within-experiment medians of positive log10 CFU/g. Correlation p-values were unadjusted. Late-sample analyses were conditional on an available fecal measurement; **Figure 2F** displays this late sampling window.

### GC-MS culture, extraction, and acquisition

ST1-75, ST1-68, ST1-6, VPI10463, and R20291 were grown for six hours in chemically defined *C. difficile* minimal medium (CDMM). Each strain was represented in three experimental batches, with an uninoculated medium sample processed in parallel within each batch. Cultures were centrifuged at 4,000 × g for 10 minutes at 4 °C to separate cells from extracellular medium. For extracellular samples, 8 microliters of medium was added to 1 mL methanol pre-cooled to −80 °C. Cell pellets were extracted in 1 mL methanol pre-cooled to −80 °C and sonicated for 30 seconds using one-second pulses separated by 0.5-second pauses. Both fractions were stored at −80 °C for 24 hours, thawed, and centrifuged at 13,000 × g for 10 minutes at room temperature. Supernatants were collected and dried for six hours in a vacuum centrifuge.

Dried extracts were resuspended in 50 microliters methoxyamine (Thermo Fisher Scientific, TS-45 840) and shaken at 1,400 rpm for two hours at 30 °C. After addition of 80 microliters N-methyl-N-(trimethylsilyl)trifluoroacetamide containing 1% trimethylchlorosilane (Thermo Fisher Scientific, TS-48 815), samples were transferred to autosampler vials with 70 microliters ethyl acetate and incubated for 30 minutes at 37 °C. Derivatized metabolites were analyzed on an Agilent 8890 gas chromatograph coupled to a 5977B mass-selective detector. One microliter was injected and samples were acquired at gain factor 5. The oven was held at 60 °C for one minute, increased at 7.3 °C per minute to 250 °C, and held at 250 °C for 10.5 minutes. Helium carrier-gas flow was maintained at 1 mL per minute. Targeted metabolite signals were processed using Agilent MassHunter Qualitative Analysis 10.0 and Quantitative Analysis B.08.00.

The intracellular dataset contained processed peak signals for blank medium and the five strains. For each bacterial sample, the signal from the blank medium in the same experimental batch was subtracted metabolite by metabolite. The resulting five-strain by three-batch design was evaluated across all 18 measured metabolites.

Metabolites were screened for signal above matched background using a one-sided test of the blank-corrected grand mean while accounting for additive strain and batch effects. Interaction/residual variation supplied eight residual degrees of freedom. Benjamini-Hochberg correction was applied across 18 metabolites, and 13 metabolites with positive corrected means at q < 0.05 were retained for visualization. Retained features were standardized, strain identity was represented with five dummy variables, and a two-component PLS regression was used for PLS-DA. Samples, strain centroids, and scaled X-loading vectors were overlaid in the same plot. This was a descriptive projection of the full dataset; classifier performance was not estimated.

The extracellular dataset contained processed peak signals for blank medium and the same five strains: ST1-75, ST1-68, ST1-6, VPI10463, and R20291. Raw signals were retained without normalization to the total signal in each sample. For each bacterial sample, the extracellular value was compared with the blank medium from the same experimental batch. Matched log2 ratios of bacterial-sample signal to blank-medium signal were used to display endpoint residual signal. Feature-specific half-minimum replacement was applied only when a zero required a logarithmic display or test; raw values without replacement were retained for comparisons with the range observed in blank medium. Thirteen metabolites were continuously quantified, defined as positive in at least two of three samples in every group, including blank medium.

For each of these 13 extracellular metabolites, global strain differences were assessed using a pseudo-F statistic accounting for additive strain and batch effects, with 9,999 permutations of strain labels restricted within experimental batches. Benjamini–Hochberg correction was applied across the 13 global tests. ST1-75 and ST1-68 were compared using two-sided paired t-tests of their batch-matched log2 residual signals, with correction across the 15 metabolites having nonzero paired differences. The geometric residual-signal ratio was calculated as 2 raised to the mean paired log2 difference. An exact two-sided sign-flip test compared the absolute mean difference with the distribution obtained by enumerating all sign assignments across the three matched batches.

The extracellular PLS-DA used all 15 bacterial samples and the 13 continuously quantified features. For each bacterial sample from batch i, each feature was represented by its batch-matched log2 residual signal relative to the corresponding uninoculated medium sample from batch i. Features were standardized across samples, strain identity was represented with five dummy-response columns, and a two-component PLS regression was fitted using plsregress. The full-data scores, strain centroids, scaled X-loading vectors, and descriptive covariance ellipses were displayed. The projection was fitted without batch centering or other residualization and independently from the intracellular projection; the coordinates and loadings were therefore not compared numerically across compartments. No independent classification-performance estimate was obtained.

### GC-MS and BIOLOG cross-assay comparison

Seven GC-MS metabolites were mapped to explicitly named PM1 substrates: lactate, alanine, aspartate, glutamate, proline, serine, and threonine. Because GC-MS labels did not encode stereochemistry, the L-form PM1 substrate was used as the primary mapping and an any-stereoisomer mapping was retained as a sensitivity analysis. We tested whether strain– substrate pairs with positive PM1 calls showed deeper extracellular depletion by permuting positive calls among the four strains measured by both assays—ST1-75, ST1-68, ST1-6, and VPI10463—within each substrate (9,999 permutations). Direct correspondence was not detected for the primary mapping (p = 0.4585) or the any-stereoisomer sensitivity (p = 0.3188).

Whole-profile correspondence was assessed across ST1-75, ST1-68, ST1-6, and VPI10463. GC-MS distances were calculated from feature-standardized mean matched log-ratios across the 13 continuously quantified metabolites, and PM1 distances were calculated using Jaccard distance across all 95 binary calls. An exact Mantel-style comparison over strain-label permutations gave Spearman ρ = −0.0857 and p = 0.875.

## BIOLOG PM1 call generation

BIOLOG Phenotype MicroArray PM1 plates contain a negative-control well and 95 wells that each provide a different carbon-source environment. Assays were conducted under anaerobic conditions and recorded as absorbance time courses. Substrate-response profiles were generated for 21 ST1 strains and VPI10463 from 30 plates: three plates each for ST1-12, ST1-68, ST1-75, and VPI10463 and one plate for each other strain. Analyses used the PM1 substrate panel. Measurements collected before 24 hours were retained. Each well was baseline-corrected to its first observation, and negative baseline-corrected values were set to zero. Separately for each plate, the MATLAB workflow modeled baseline-corrected absorbance using time, substrate, their interaction, and a well-specific random slope for time, with the negative-control well as the reference. A substrate was called positive if it appeared in any non-intercept fixed-effect coefficient with a positive estimate and p ≤ 0.05/95. The implemented rule therefore includes significant positive substrate main effects and time-by-substrate interactions. For ST1-12, ST1-68, ST1-75, and VPI10463, a substrate was positive in the final strain profile if any of three plates was positive; other strains had one plate. All 30 plates showed the required positive response to alpha-D-glucose and no positive response in the negative-control well. No additional plate-exclusion or replicate-concordance rule was applied.

The workflow produced a matrix of 22 strains by 96 wells, including the negative-control well. Removing that well left 95 substrates and 700 positive calls. For every ST1/VPI10463 pair, nutrient overlap was the fraction of VPI10463-positive substrates also positive for the ST1 candidate. Additional resource access was the number of candidate-positive substrates outside that overlap. Total relative access, ΔB, was candidate total breadth minus VPI10463 total breadth.

Because VPI10463 was fixed across comparisons and had 23 total positive calls, ΔB = B_ST1_ - 23. Amino-acid breadth was the unweighted sum across the 16 environments annotated as L-aspartic acid, L-proline, D-alanine, D-serine, L-glutamic acid, L-asparagine, D-aspartic acid, L-glutamine, Gly-Asp, D-threonine, Gly-Glu, L-serine, L-threonine, L-alanine, Ala-Gly, and Gly-Pro. VPI10463 had three positive calls in this class, so ΔB_AA_ = B_ST1,AA_ - 3. Subtracting either fixed VPI10463 value changes the ecological interpretation and intercept but not rank correlations or fitted resource slopes. R20291 did not have a PM1 profile in this dataset and was therefore retained in GC-MS analyses but excluded from direct cross-assay comparisons.

The VPI10463-rooted, recombination-filtered tree determined the order of the 21 ST1 strains in **Figure 4**. Weakly supported internal branches were collapsed for display as described below; patristic distances and phylogenetic tests used the full fitted tree. PM1 substrates were grouped by chemical class, retaining plate order within each class.

We tested the association between ΔBAA and the co-infection protection score using Spearman correlation. Because VPI10463 contributed the same three-call constant to every comparison, this correlation is identical to that for ST1 amino-acid breadth. The protection score came from the composite disease-course analysis in which zero was assigned only when weight was missing at a recorded terminal event. A 5,000-resample strain bootstrap with random seed 38 provided the percentile interval. The dashed ordinary least-squares line in **Figure 4D** was used as a visual guide.

For the disease-adjusted display, tied ranks of ΔBAA and protection were each residualized against an intercept and the tied rank of the mono-colonization disease score; Pearson correlation between the two residual vectors gives the partial-rank statistic shown in **Figure 4E** and **Supplementary Figure S2B,C**, with a two-sided t approximation using 18 degrees of freedom to account for the disease covariate. To assess unequal plate replication, we repeated the associations using one PM1 plate per strain across all 81 combinations of the three plates available for ST1-12, ST1-68, ST1-75, and VPI10463, retaining the single plate available for each other strain. These yielded nine distinct rank-predictor patterns; we report the full effect-size ranges. We then compared protection models with disease alone and with disease plus ΔBAA. Ordinary least squares supplied adjusted R2 and leave-one-strain-out predictions. A covariance-aware generalized least-squares analysis used the covariance among the 21 protection-score coefficients estimated from the primary mixed-effects model, plus a fitted residual-variance component. Model comparison used AICc and a one-degree-of-freedom likelihood-ratio test. A reduced-model residual-permutation test used 10,000 permutations to test the increment attributable to amino-acid access. These analyses evaluate explanatory fit; leave-one-strain-out RMSE was used to assess whether that increment produced held-out predictive improvement.

Chemical-class breadths were calculated for carbohydrates, carboxylic acids, amino acids, fatty acids, alcohols, amines, amides, and esters. Spearman correlations with protection were corrected across the eight classes using the Benjamini–Hochberg procedure. **Supplementary Figure S2D** displays the unadjusted coefficients as bars and uses asterisks only for classes with q < 0.05. The eight-class comparison was exploratory; direct examination of the complete amino-acid class was motivated by the previous ST1-75 competition study and the faster proline consumption by ST1-75 than ST1-68. Specificity analyses compared the 16-well amino-acid score with breadth across the other 79 wells and fitted a joint model containing disease, amino-acid breadth, and non-amino-acid breadth. Robustness was assessed by removing each amino-acid-class well in turn and by leave-one-strain-out influence analysis. **Supplementary Figure S2B,C** displays the specificity and well-deletion analyses; complete strain-influence and held-out prediction results are retained in the analysis outputs.

### Protein-family and nutrient-overlap analyses

We used global protein-family (PGFam) assignments in feature exports from the Bacterial and Viral Bioinformatics Resource Center (BV-BRC) for the 21 ST1 isolates and VPI10463 to calculate pathogen-centered overlap (Spragge et al. 2023). Distinct PGFams assigned to coding sequences were counted once per genome; unassigned sequences were omitted and assigned hypothetical proteins were retained. Overlap was the fraction of distinct VPI10463 PGFams also present in the candidate. The VPI10463 annotation was genome 1496.1555, corresponding to assembly GCA_001 995 155.1; we also used this assembly for outgroup sensitivity, separately from the primary phylogenetic outgroup. Strain-to-genome mappings are listed in **Supplementary Table S1**.

We calculated nutrient overlap across all PM1 substrates, the amino-acid class, and the remaining substrates, using VPI10463-positive denominators of 23, 3, and 20, respectively. Across all 21 strains, raw and disease-adjusted Spearman associations used average ranks; adjusted tests residualized predictor and protection ranks against an intercept and disease rank, with 18 degrees of freedom. Percentile intervals used 5,000 strain-level bootstraps with seed 20 261 001, and ranks were recomputed in each draw. One constant amino-acid-overlap draw was omitted from its raw and adjusted intervals. We retained complete leave-one-strain-out results and repeated nutrient-overlap tests across all 81 one-plate-per-strain combinations, recalculating VPI10463 denominators for each selection. Benjamini–Hochberg correction treated the six nutrient-overlap tests as one family and three protein-family tests as a separate family: raw overlap, disease-adjusted overlap, and relative amino-acid access adjusted for disease and protein-family overlap (17 degrees of freedom). These exploratory analyses used strain-level score estimates; the covariance-aware and phylogeny-adjusted analyses of relative amino-acid access are described separately.

### Consumer-resource model

The consumer-resource system assumes binary substrate access; equal supply, uptake, and conversion for every resource class; no cross-feeding; no spatial structure; and no measured resource concentrations. For each ST1/VPI10463 pair, O denotes the number of overlapping resources, BST1 the candidate’s total resource access, and BVPI the pathogen’s total access. Access outside the overlap is BST1 minus O for the candidate and BVPI minus O for the pathogen. The aggregate limitation parameter λ combines loss, dilution, and supply limitation on the count scale. Because the assay does not estimate the kinetic parameters needed to fit λ, we evaluated 501 evenly spaced values from 0 to 82, the maximum observed breadth.

Rare-invasion scores indicate whether a strain can increase from low abundance in the presence of the established competitor; positive scores indicate invasion. The rare-invasion scores were calculated as follows:

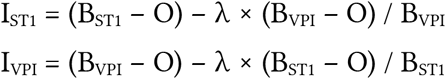

At each value of λ, we first assessed whether each strain could persist alone: its total resource access had to exceed λ. If only one strain met this condition, the outcome was exclusion of the other strain; if neither did, neither strain was feasible. Only when both strains were feasible did the signs of IST1 and IVPI determine the outcome: two positive scores indicated coexistence, one positive score indicated exclusion of the other strain, and two nonpositive scores indicated priority effects. Equality to zero was treated as non-invasion. ST1 persistence was the fraction of the grid for which ST1 either coexisted with or excluded VPI10463. The upper part of **Figure 4F** displays VPI10463-centered overlap and additional candidate resource access, and the lower part displays the corresponding outcome fractions for ST1-75, ST1-6, and ST1-68. The persistence fraction was used as a theory-based sensitivity summary of the binary call structure, not as a calibrated *in vivo* kinetic estimate.

### Adaptive-immunity and immune-cell analyses

The RAG1 challenge trajectories in **Figure 5A** show all observed values for ten mice per group; thin lines represent individual mice and bold lines represent group means among observed animals. Nine previously uncolonized wild-type mice died or were removed at a humane endpoint before the scheduled final day. Their final observed weights are marked with grey crosses, and subsequent values remain missing; no terminal weight was imputed. The model effects in **Figure 5B** were estimated from observed relative weights, with condition as a fixed effect and random intercepts for day and animal, matching the disease-course model form used for the main mouse comparisons. The RAG1 analysis retained observed relative weights without adding terminal-event score zeros. Each coefficient compares one ST1-75-colonized arm with the same previously uncolonized wild-type reference.

In a separate experiment, we compared colonic lamina-propria immune-cell populations in ST1-75-colonized mice and uncolonized controls before VPI10463 challenge (three mice per group; **Figure 5C,D**). Colonized mice received two antibiotic courses; controls received one or two. Cells were isolated from the entire colon and analyzed on an LSR II flow cytometer, collecting 100,000 events per sample for each staining panel. The adaptive panel summarized B cells, CD19-negative/TCRβ-negative cells, natural killer cells, T cells, CD4-positive cells, and CD8-positive cells. The innate panel summarized dendritic cells, CD11b-positive cells, lymphoid and myeloid dendritic cells, macrophages, monocytes, and neutrophils. Within each staining panel, events were retained if all scatter and fluorescence measurements lay within the per-channel minimum and maximum values observed among events assigned to at least one cell-type gate, pooled across the six mice. For each mouse, each cell-type fraction was its gate-positive count divided by the total retained-event count in that panel. Fractions were compared between groups using Wilcoxon rank-sum tests.

### Residual-microbiome and genomic analyses

DNA was extracted from fecal samples using a previously described laboratory protocol (Taur et al. 2018). The V4–V5 region of the 16S rRNA gene was amplified with Q5 High-Fidelity DNA polymerase (New England Biolabs, M0491) in 25-microliter reactions containing 5 microliters Q5 buffer, 5 microliters Q5 enhancer, 0.5 microliters 10 mM dNTPs, 0.625 microliters Q5 polymerase, 2.5 microliters each of 10 micromolar indexed forward and reverse primers, and 1 ng template DNA, with water added to the final volume. Cycling conditions were 96 °C for 10 minutes; 35 cycles of 96 °C for 10 seconds, 51 °C for 30 seconds, and 72 °C for 30 seconds; and a final extension at 72 °C for five minutes. Amplification was confirmed by agarose-gel electrophoresis. Products were purified with AxyPrep magnetic beads, quantified using the Quant-iT PicoGreen double-stranded DNA assay (Invitrogen, P7589), normalized, and pooled. Sequencing libraries were prepared with the KAPA LTP Library Preparation kit (07 961 871 001), quantified by PicoGreen, normalized, and sequenced with an Illumina MiSeq Reagent Kit v3. Sequence processing followed the previously described laboratory workflow (Liao et al. 2024).

Residual-microbiome analysis began from the resulting genus-level abundance table. *Clostridioides* was removed and remaining genera were renormalized within sample. Shannon diversity was compared between ST1-75 and ST1-12 at each day using Wilcoxon tests. Bray-Curtis composition was assessed with a two-group permutation test, and ten day-1 genus-level features were compared with false-discovery correction. **Figure 5E** presents the Shannon-diversity summary; **Supplementary Figure S3A,B** shows the Bray-Curtis and day-1 genus-level comparisons.

Whole-genome phylogenetic analysis used assemblies of the 21 ST1 isolates and VPI10463/ATCC 43 255 (GCF_015 238 635.1). Public read accessions for the ST1 isolates and reference-assembly accessions are listed in **Supplementary Table S1**. Assemblies were aligned to R20291 (GCF_000 027 105.1) using NUCmer in MUMmer 4.0.1, retaining one-to-one matches of at least 95% identity and 500 bases (Marçais et al. 2018). Repeated reference regions, ambiguous or overlapping alignments, five bases flanking indels, and positions with discordant calls in an independent SKA2 alignment were excluded. Uncovered positions remained missing. Gubbins 3.4.3 identified recombination in the reference-coordinate alignment of the 21 ST1 isolates, excluding the divergent VPI10463 outgroup (Croucher et al. 2015). The union of inferred recombination intervals was removed from all genomes. Retaining sites called in every genome produced 3,133,615 positions, including invariant sites. Maximum-likelihood inference used IQ-TREE 2.4.0 with the TVM+F model selected by ModelFinder using the Bayesian information criterion, 1,000 ultrafast bootstrap replicates with nearest-neighbor interchange refinement, and 1,000 SH-aLRT replicates (Minh et al. 2020). The tree was rooted on VPI10463, which was then pruned from the display. Internal branches below 95% ultrafast bootstrap or 80% SH-aLRT support were collapsed for display only. ST1-75/ST1-68 nucleotide differences were counted across positions callable in both strains after the same quality and recombination filtering.

Pairwise patristic distances were extracted from the full fitted ST1 tree. For each trait, a two-sided Mantel-style label-permutation test used Spearman correlation between patristic distance and absolute pairwise trait difference, with 10,000 permutations in a fixed alphabetical strain order. Random seed 1 was used sequentially for total relative access, protection, and disease; seed 29 was used for amino-acid relative access. Sensitivity analyses repeated inference without recombination filtering, without the out group on the same alignment columns, and with an alternative VPI10463 assembly (GCA_001 995 155.1) on columns shared by both outgroups.

To test whether amino-acid access contributed information beyond disease potential and phylogenetic relatedness, we fitted protection scores to mono-colonization disease scores, with or without relative amino-acid access, across all 21 strains. The residual covariance combined the full sampling covariance of the protection-score estimates with independent and phylogenetic variance components. Phylogenetic covariance was calculated from branch lengths shared from the ST1 root in the full, uncollapsed recombination-filtered tree and scaled to a mean diagonal of one. Regression coefficients and nonnegative variance components were estimated by maximum likelihood separately for each model, including zero-variance boundaries.

The incremental likelihood-ratio statistic was evaluated using 1,999 parametric-bootstrap datasets simulated from the fitted disease-only model (random seed 20 260 919). Both regression and variance parameters were refitted in each simulation. The p-value was the number of simulated statistics at least as large as the observed statistic, plus one, divided by 2,000. The tree, predictor values, and estimated score-sampling covariance were held fixed. Sensitivity analyses repeated the comparison with alternative trees, collapsed unsupported branches, independent-only or phylogenetic-only residual variation, omission of score-sampling covariance, and deletion of each strain in turn.

## Data and code availability

Raw sequencing reads for the 21 ST1 isolates are available in the NCBI Sequence Read Archive under BioProject PRJNA595 724 (Dong et al. 2023). Strain-level accessions and reference genomes are listed in **Supplementary Table S1**; the assembled FASTA inputs used for phylogenetic analysis are included with the analysis code.

Experimental data, genomic FASTA inputs, and analysis code, including the protein-family and nutrient-overlap analyses, are available in the cdiff_resource_access repository at commit d740 151.

## Supplementary material

**Supplementary Figure S1.**
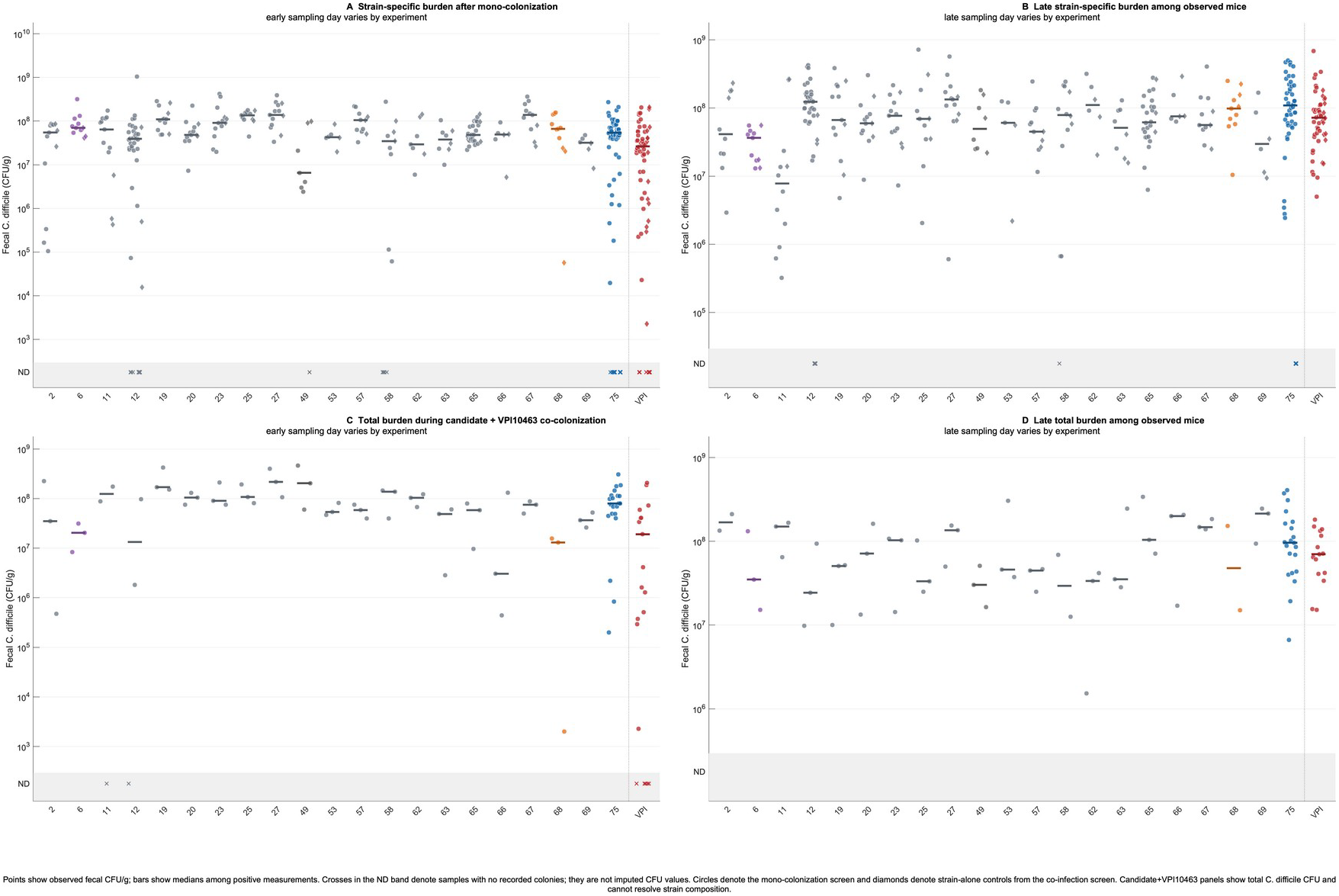
Fecal burden across the mono-colonization and co-colonization screens. (A,B) Strain-specific fecal *C. difficile* CFU/g in early and late samples after inoculation with each ST1 isolate or VPI10463 alone. Circles show measurements from the mono-colonization screen; diamonds show strain-alone controls run alongside the co-infection screen. Every ST1 isolate was recovered from at least one mouse in both windows. Positive strain-alone burdens differed among strains (early overall comparison, p = 4.19 × 10^-6^, q = 1.68 × 10^-5^; late overall comparison, p = 0.000137, q = 0.000273), but burden was not associated with mono-colonization disease (early, ρ = −0.405, p = 0.0696; late, ρ = −0.023, p = 0.921). ST1-68 was detected in all 11 early and all 11 late samples available across three experiments; its late burdens ranged from 1.04 × 10^7^ to 2.50 × 10^8^ CFU/g. (C,D) Total fecal *C. difficile* CFU/g in early and late samples from candidate+VPI10463 groups; the VPI10463 label shows VPI10463-only controls. Co-colonization plating does not resolve which strain contributed the colonies. Early total burdens did not differ detectably among candidate groups (p = q = 0.0627) and were not associated with protection (ρ = 0.265, p = 0.245). Late total burdens differed among the mice with an available sample (p = 0.0335, q = 0.0446) but did not track protection (ρ = −0.119, p = 0.605). Points show observed values and horizontal bars show medians among positive measurements. Crosses in the non-detect (ND) band indicate that no colonies were recorded; they are not imputed CFU values. Early samples were collected on experiment-specific day 1 or 2 and late samples on day 6 or 7. Late panels include only mice with an observed fecal measurement and are therefore conditioned by death, humane-endpoint removal, and other missingness. ST1-75, ST1-68, and ST1-6 are highlighted in blue, orange, and purple, respectively; the remaining ST1 isolates are shown in grey.

**Supplementary Figure S2.**
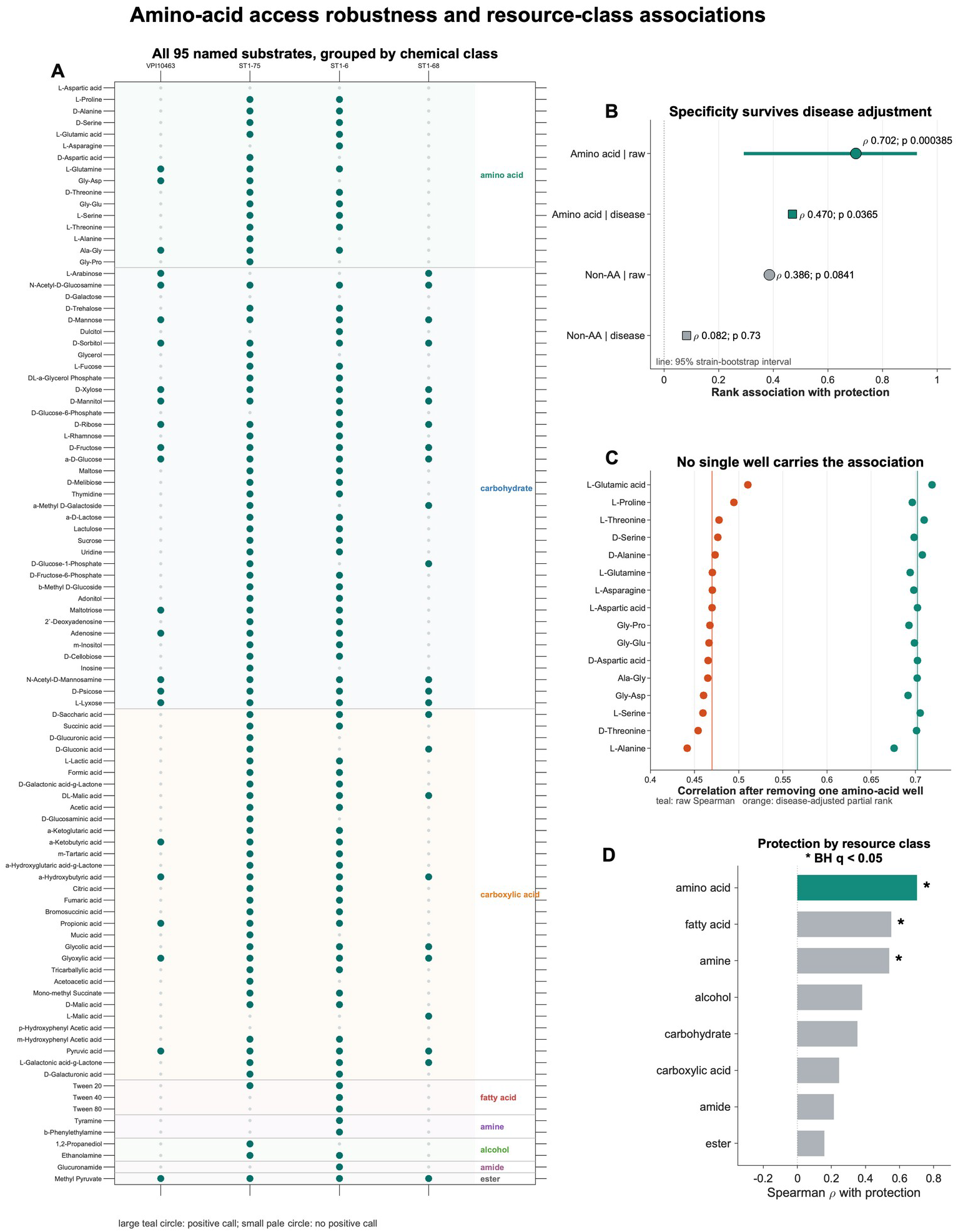
Amino-acid access robustness and resource-class associations. (A) Positive PM1 calls for VPI10463, ST1-75, ST1-6, and ST1-68 across all 95 named substrates, grouped by chemical class and retaining PM1 plate order within class. Large teal circles indicate positive binary response calls under the workflow described in Methods; small pale circles indicate no positive call. (B) Raw and disease-adjusted rank associations of protection with amino-acid breadth and breadth across the remaining 78 PM1 environments. The horizontal line through the raw amino-acid estimate is its 5,000-resample strain-bootstrap 95% interval. Because VPI10463 is fixed, subtracting its class breadth gives the same rank statistics.(C) Raw and disease-adjusted correlations after removing each amino-acid-class well from the unweighted score one at a time. Vertical lines mark the complete 16-well values. (D) Spearman associations between chemical-class breadth and protection. Asterisks denote Benjamini– Hochberg q < 0.05 across the eight class comparisons. Amino acids are highlighted in teal.

**Supplementary Figure S3.**
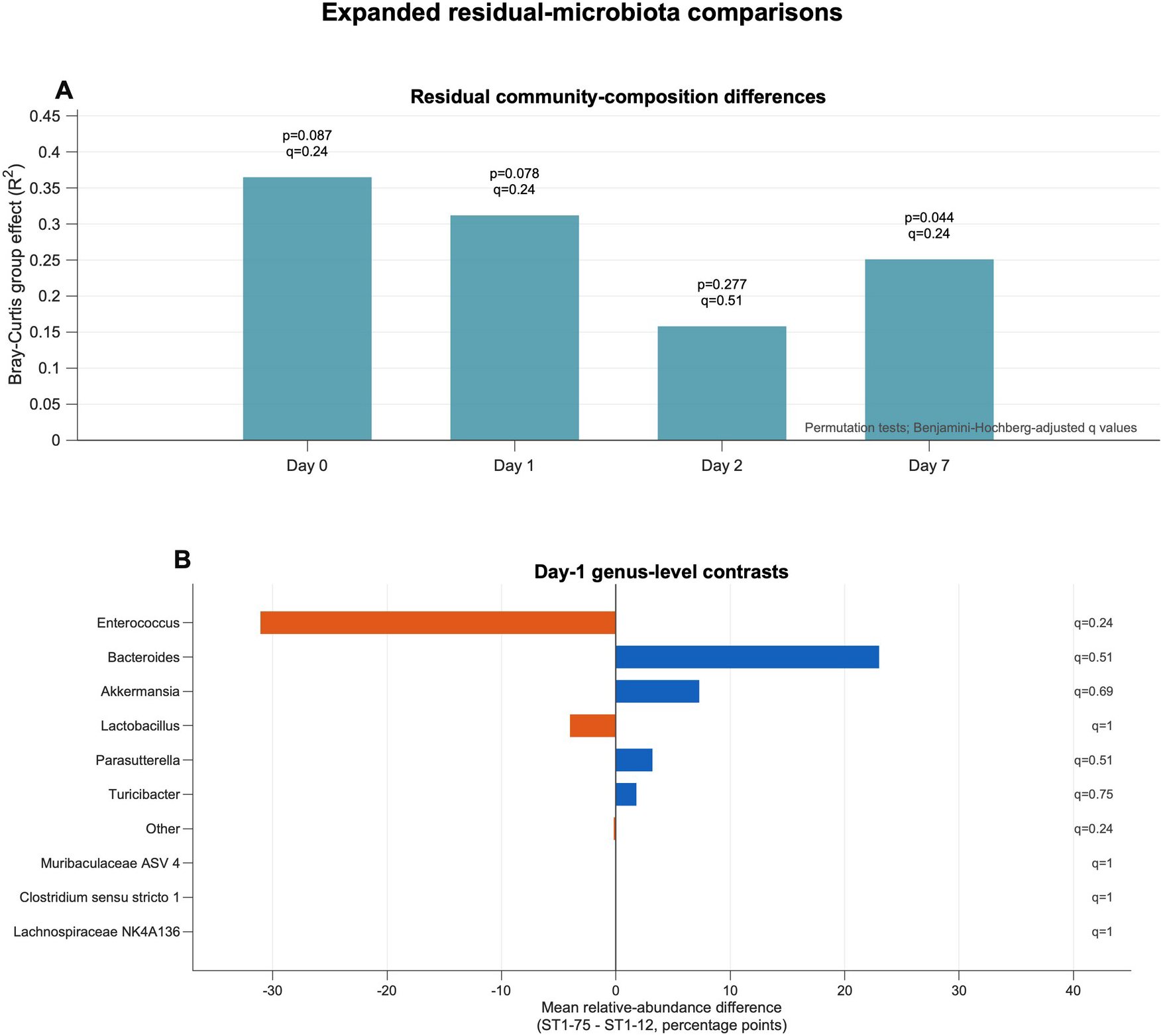
Expanded residual-microbiota comparisons. (A) Bray-Curtis group-effect sizes comparing residual microbiota after colonization with ST1-75 or ST1-12 on days 0, 1, 2, and 7, following removal of *Clostridioides* and renormalization of the remaining genus-level community. Each strain and day contains 3–5 mice. Bars show the permutation-test R^2^ values; labels give the corresponding unadjusted p and adjusted q values. (B) Day-1 differences in mean relative abundance for the ten genus-level features examined. Positive values indicate greater mean abundance after ST1-75 colonization and negative values indicate greater mean abundance after ST1-12 colonization; labels give adjusted q values. Each group contained five mice. None of the community-composition or genus-level comparisons survived false-discovery correction.

**Supplementary Table S1.** Public sequencing accessions, reference genomes, and BV-BRC annotation identifiers. Raw reads for the 21 clinical ST1 isolates are deposited under BioProject PRJNA595 724 (Dong et al. 2023). ST1 entries identify raw-read records; reference-genome entries identify assembled sequences. The exact assembled FASTA inputs used in the phylogenetic analysis and the BV-BRC feature exports used for protein-family analysis are included with the analysis code.

| Strain | BioSample | SRA run | BV-BRC genome ID |
| --- | --- | --- | --- |
| ST1-2 | SAMN13 566 372 | SRR10 693 941 | 1485.318 |
| ST1-6 | SAMN13 566 324 | SRR10 693 997 | 1485.330 |
| ST1-11 | SAMN13 566 371 | SRR10 693 942 | 1485.310 |
| ST1-12 | SAMN13 566 370 | SRR10 693 943 | 1485.309 |
| ST1-19 | SAMN13 566 326 | SRR10 693 994 | 1485.315 |
| ST1-20 | SAMN13 566 335 | SRR10 693 981 | 1485.316 |
| ST1-23 | SAMN13 566 336 | SRR10 693 980 | 1485.314 |
| ST1-25 | SAMN13 566 328 | SRR10 693 989 | 1485.317 |
| ST1-27 | SAMN13 566 330 | SRR10 693 987 | 1485.335 |
| ST1-49 | SAMN13 566 378 | SRR10 693 934 | 1485.322 |
| ST1-53 | SAMN13 566 406 | SRR10 693 902 | 1485.324 |
| ST1-57 | SAMN13 566 317 | SRR10 694 004 | 1485.321 |
| ST1-58 | SAMN13 566 423 | SRR10 693 884 | 1485.325 |
| ST1-62 | SAMN13 566 360 | SRR10 693 954 | 1485.326 |
| ST1-63 | SAMN13 566 320 | SRR10 694 001 | 1485.329 |
| ST1-65 | SAMN13 566 318 | SRR10 694 003 | 1485.327 |
| ST1-66 | SAMN13 566 292 | SRR10 694 159 | 1485.332 |
| ST1-67 | SAMN13 566 325 | SRR10 693 995 | 1485.328 |
| ST1-68 | SAMN13 566 403 | SRR10 693 907 | 1485.333 |
| ST1-69 | SAMN13 566 435 | SRR10 693 846 | 1485.331 |
| ST1-75 | SAMN13 566 366 | SRR10 693 947 | 1485.334 |

Reference genome assemblies
| Strain | Assembly accession | BioSample | Use |
| --- | --- | --- | --- |
| R20 291 | GCF_000 027 105.1 | SAMEA3 138 243 | Reference coordinates<br>for genome alignment |
| VPI10 463 (ATCC<br>43 255) | GCF_015 238 635.1 | SAMN13 293 454 | Primary phylogenetic<br>outgroup |
| VPI10 463 | GCA_001 995 155.1 | SAMN06 215 604 | Alternative outgroup<br>for sensitivity analysis;<br>PGFam annotation<br>genome 1496.1555 |

## Notes

### Competing Interest Statement

The authors have declared no competing interest.

### Funding

- National Institute of Allergy and Infectious Diseases: R01 AI196346, P01 AI179406

