## Supplementary figures and images for "Resource-access breadth marks protective strains against toxigenic *Clostridioides difficile*"

### Supplementary Figure S1

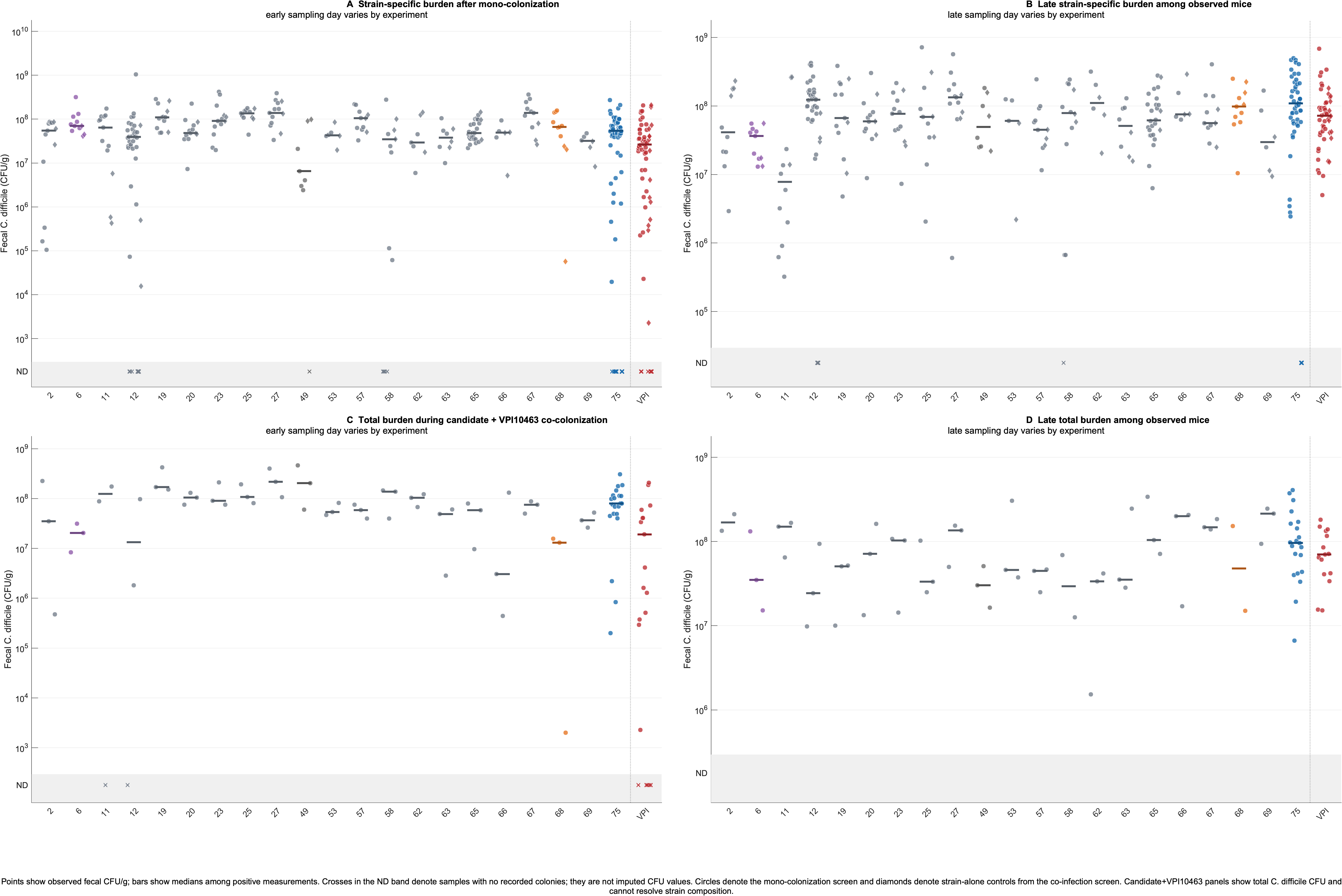

### Supplementary Figure S2

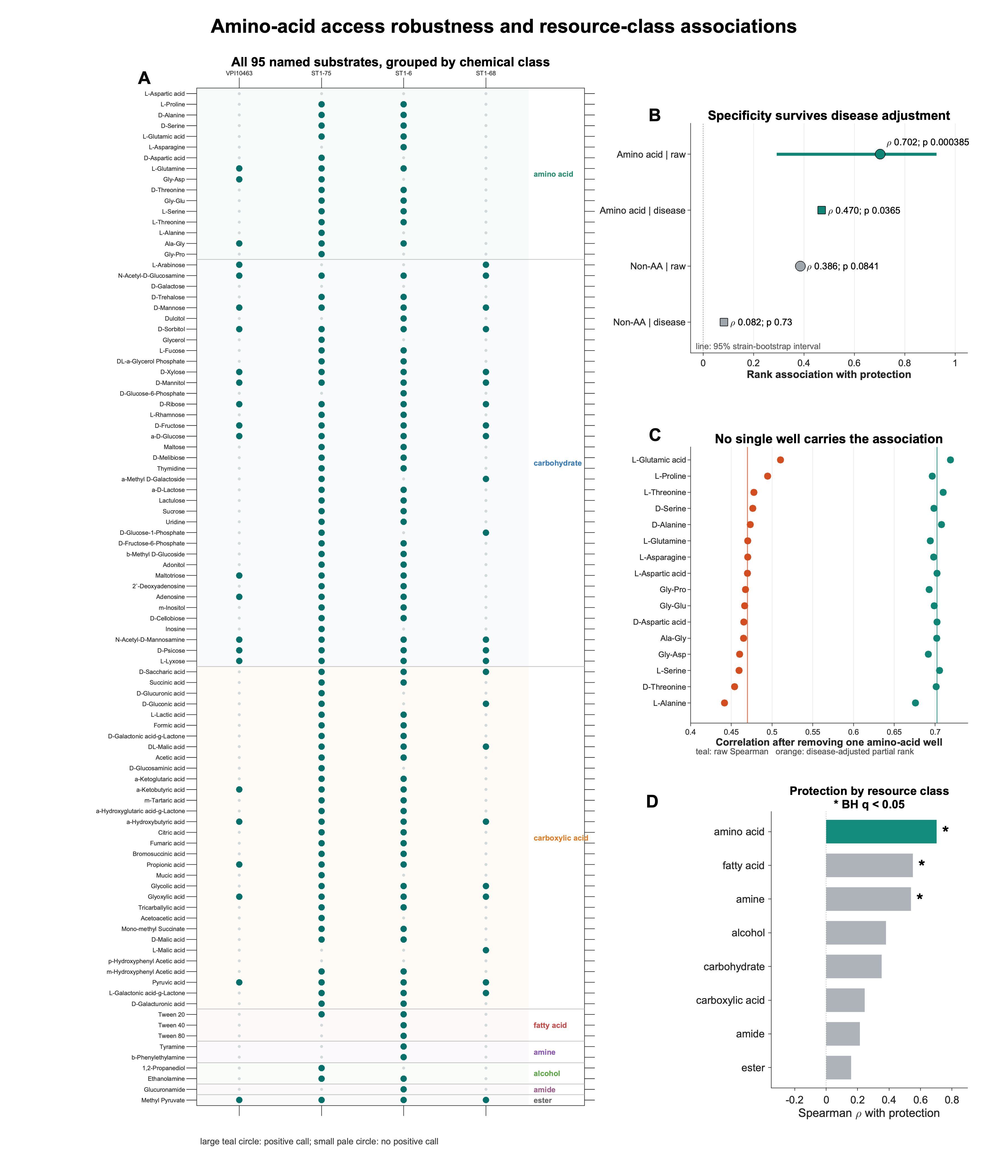

### Supplementary Figure S3

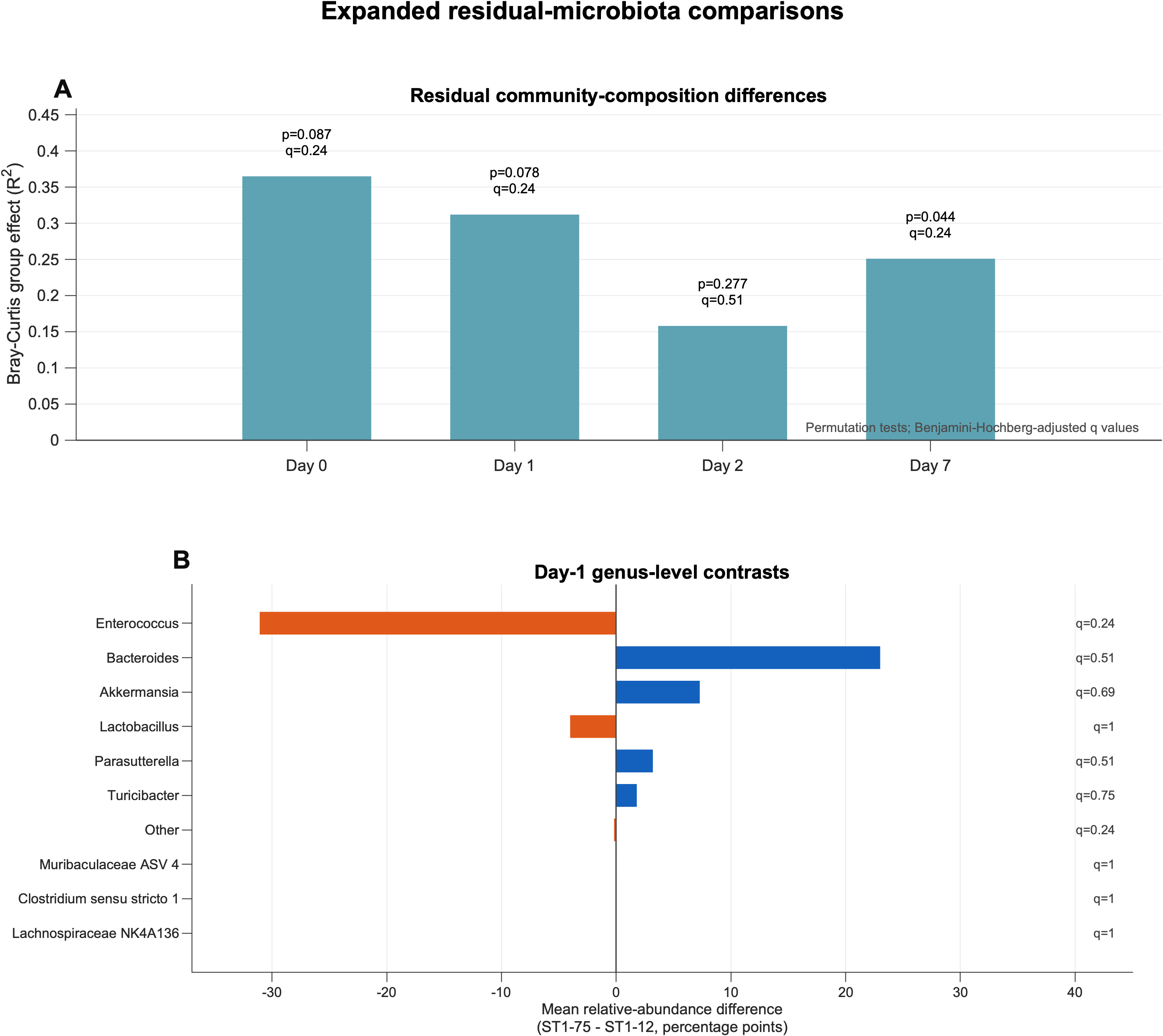
